# Nuclear polysaccharides maintain H3K9me3-heterochromatin and genomic stability

**DOI:** 10.1101/2025.08.12.669888

**Authors:** Xiuxiao Tang, Ranran Dai, Li Qing, Zhida Zhang, Lizi Lu, Hancheng Lin, Wei Dan, Yuqi He, Xinyi Liu, Wakam Chang, Yang Mao, Shisheng Sun, Junjun Ding

## Abstract

Polysaccharides are known to be synthesized by enzymes in the endoplasmic reticulum and Golgi apparatus and transported through the secretory pathway to the cell surface or extracellular space^1^, where they mediate essential biological processes^2-6^. While classical localization and functions of polysaccharides are well established, their presence and potential roles in the nucleus remain unclear. Here, we demonstrate that *N*-glycans, a type of polysaccharides, are present in the cell nucleus and modify inner nuclear membrane (INM) proteins across diverse cell types—a modification referred to as *N*-linked glycosylation (*N*-glycosylation). *N*-glycosylation is enriched in chromatin regions marked by H3K9me3 and long interspersed nuclear element-1 (LINE-1) retrotransposons. *N*-glycosylation inhibition and INM protein *N*-glycosylation site mutation both downregulate H3K9me3 within lamina-associated domains (LADs) and lead to genomic instability. Mechanistically, *N*-glycosylation regulates the interaction between the histone H3K9 methyltransferase SETDB1 and INM proteins, promotes the association of SETDB1 with the INM, and maintains H3K9me3. Moreover, we reveal that canonical *N*-glycan biosynthetic machinery in ER contributes to the *N*-glycosylation of INM proteins. These findings uncover a previously unrecognized nuclear role for polysaccharides, broadening our understanding beyond their traditional subcellular distributions and functional profiles.

---

Saccharides, one of the four major classes of biomolecules, can covalently modify proteins, lipids, and RNAs to form glycoproteins, glycolipids, and glycoRNAs, respectively ^1,2^. Monosaccharides serve as the basic building blocks of saccharides; they can be attached individually to a protein or further extended by the addition of more monosaccharides to form polysaccharides ^1^. These polysaccharides chains are referred to as glycans ^3^.

Different classes of saccharides exhibit specific subcellular localizations. Among them, only the monosaccharide *N*-acetylglucosamine (GlcNAc)—a glucose derivative in which the nitrogen atom of the amino group (–NH₂) is acetylated—has been shown to modify nuclear proteins as a single unit within the nucleus, a phenomenon known as *O*-linked *N*-acetylglucosamine modification (*O*-GlcNAcylation) ^3^. In contrast, polysaccharides modification relies on enzymes in the endoplasmic reticulum and Golgi apparatus, following a secretory pathway that delivers polysaccharides to the cell membrane or extracellular space ^3-5^. Polysaccharides are present in nearly all eukaryotic organisms and play key roles in biological processes at the cell membrane, such as signal transduction, immune responses, and cell adhesion ^2,6-8^. Their dysregulation is associated with various diseases, including autoimmune disorders and cancers ^9-12^. However, despite extensive studies on the functions of polysaccharides at the cell membrane, their presence and potential roles in the nucleus remain unknown.

Here, we show that *N*-glycans—a class of polysaccharides covalently linked to the nitrogen atom (*N*) of asparagine residues—are present in the nucleus and modify inner nuclear membrane (INM) proteins across various cell types, as revealed by StrucGP software for site-specific glycoproteomic interpretation ^13^ as well as further validation using immunofluorescence and our established GlycoChIP-seq approach. Using global *N*-glycosylation inhibition and *N*-glycosylation site (*N*-glycosite) mutation on INM protein, we further confirm that *N*-glycosylation could maintain H3K9me3-heterochromatin and genomic stability. Furthermore, we demonstrate that canonical *N*-glycan biosynthetic machinery in ER contributes to the *N*-glycosylation of INM proteins. Together, these findings highlight a nuclear role for polysaccharides and broaden their known regulatory functions.

## Inner nuclear membrane proteins can be *N*-glycosylated across diverse cell types

To explore the potential existence of polysaccharides in the nucleus, we isolated nuclei from five cell types with different species origins and developmental potentials. These included mouse totipotent-like stem cells (TLSCs), capable of giving rise to both embryonic and extraembryonic tissues; mouse and human embryonic stem cells (ESCs), which can differentiate into all three germ layers; mouse pre-induced pluripotent stem cells (pre-iPSCs), with lower potency compared to ESCs; and mouse embryonic fibroblasts (MEFs), representing fully differentiated somatic cells (Fig. 1a) ^14,15^. We performed structural and site-specific glycoproteome analyses on these isolated nuclei using the StrucGP software ^13^, and found *N*-glycosylation on inner nuclear membrane (INM) proteins SUN1, SUN2, TMPO, and LEMD3 across all five cell types. High-quality mass spectrometry data further confirmed that these glycoproteins were modified specifically by high-mannose *N*-glycans (Fig. 1a, b and Extended Data Fig. 1).

**Fig. 1.**
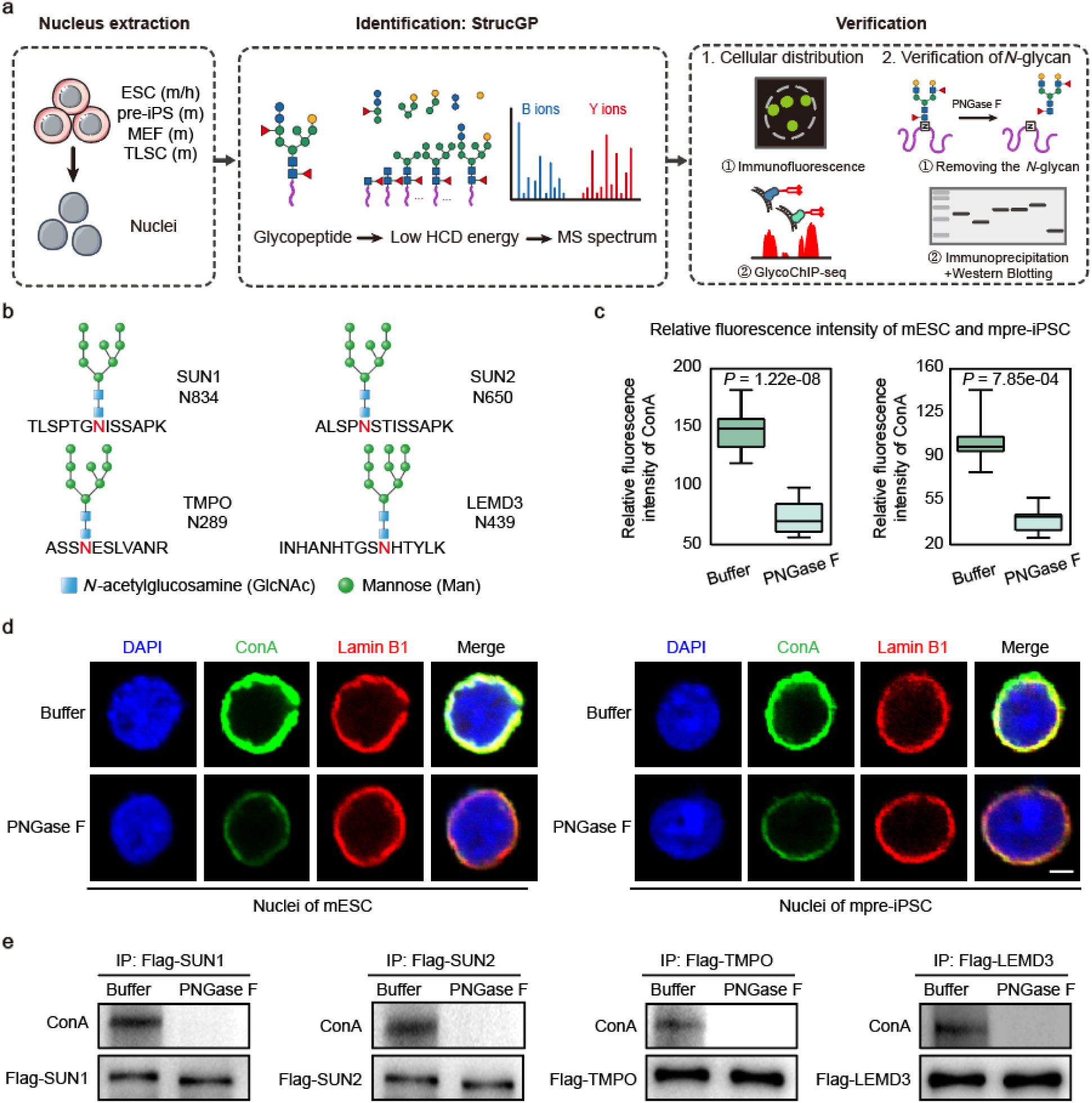
Inner nuclear membrane proteins can be *N*-glycosylated across diverse cell types. **a**, Schematic of the identification and verification of *N*-glycosylation in the nucleus, m indicates mouse, h indicates human. **b**, Site-specific *N*-glycan structure analysis of the *N*-glycosylated INM proteins identified in five cell types shown in **a**. **c**, The quantitative analysis of fluorescence signals of ConA in **d**, *P*-values by two-tailed t-test. **d**, Immunofluorescence of cell nuclei before and after treatment with PNGase F, scale bar is 5 μm. **e**, Blotting of purified INM proteins before and after treatment with PNGase F, these experiments were performed three times, with similar results.

We further confirmed the presence of *N*-glycosylation in the nucleus and the *N*-glycosylated states of INM proteins using two sets of experiments (Fig. 1a). The nuclear localization of *N*-glycosylation was primarily assessed by immunofluorescence and our established GlycoChIP-seq method (Fig. 1a and Fig. 2). Specifically, isolated nuclei were stained with ConA, a lectin that selectively recognizes high-mannose *N*-glycans ^16^. As the nuclear lamina lies directly beneath and associates closely with the INM, we used Lamin B1, a core lamina component in mESCs that interacts with all identified glycoproteins, as an INM marker ^17^. ConA signals strongly colocalized with Lamin B1 in isolated nuclei and were markedly reduced by PNGase F, which removes *N*-glycans from glycoproteins ^2^ (Fig. 1c, d), while glucose supplementation—the substrate for *N*-glycosylation—increased the signals (Extended Data Fig. 2a, b). These results demonstrated that *N*-glycosylation was mainly localized to the inner nuclear membrane region.

**Fig. 2.**
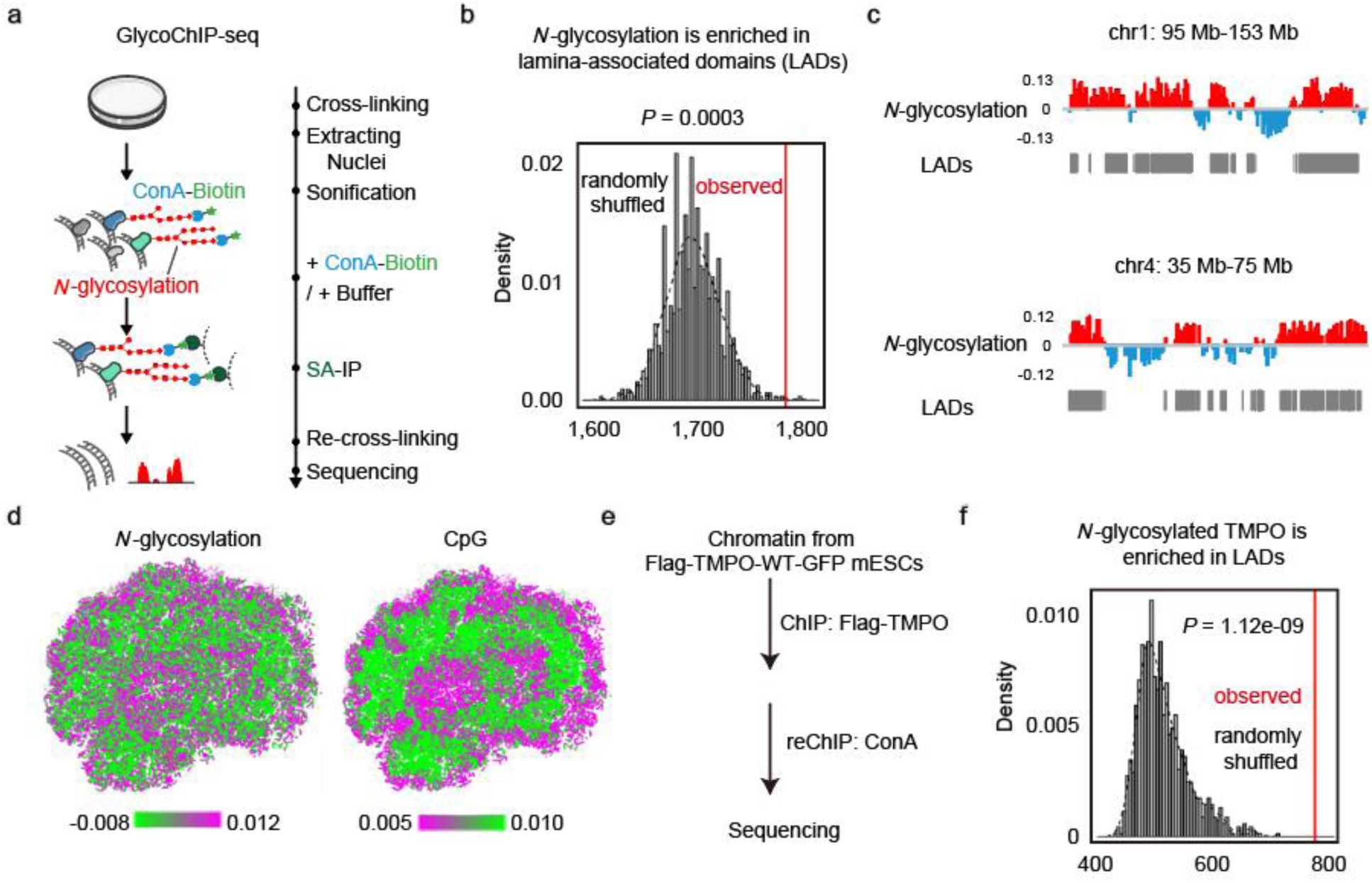
*N*-glycosylation and *N*-glycosylated INM protein are enriched in chromatin. **a**, Schematic of GlycoChIP-seq to detect the chromatin binding of *N*-glycosylation. **b**, Permutation test showing *N*-glycosylation is enriched in lamina-associated domains (LADs), *P*-value by one-tailed test based on the assumption of normal distribution. **c**, Representative genome tracks of buffer-normalized *N*-glycosylation GlycoChIP-seq from mESCs. Gray bars indicate regions defined as LADs. **d**, Representative 3D genome structure of mESCs, colored by *N*-glycosylation frequency (left) and CpG frequency (right), cell was chosen randomly. **e**, Schematic of serial ChIP-seq to detect the chromatin binding of *N*-glycosylated TMPO. **f**, Permutation test showing *N*-glycosylated TMPO is enriched in LADs, *P*-value by one-tailed test based on the assumption of normal distribution.

To confirm the *N*-glycosylation of INM proteins, we enriched glycoproteins from nuclear proteins using ConA under denaturing conditions to disrupt protein-protein interactions, and probed the above identified INM proteins using Western blotting. All of the above INM proteins were present in the ConA-enriched *N*-glycosylated nuclear proteome (Extended Data Fig. 2c). The *N*-glycosylation of INM proteins was further confirmed by purifying them and then detecting their *N*-glycosylated states using a lectin blot with ConA (Extended Data Fig. 2d). After treating the purified INM proteins with PNGase F, the *N*-glycosylation signals disappeared from ConA lectin blot (Fig. 1e).

Taken together, all these results confirm that INM proteins can be *N*-glycosylated and are mainly modified by high-mannose *N*-glycans.

## *N*-glycosylation and *N*-glycosylated INM protein are enriched in chromatin

In light of the association between the inner nuclear membrane and chromatin ^18^, and to further substantiate the nuclear localization of *N*-glycosylation as well as explore its potential binding to chromatin, we developed a lectin-based genome-wide chromatin immunoprecipitation followed by sequencing (GlycoChIP-seq) (Fig. 2a). GlycoChIP-seq utilizes ConA-biotin to recognize *N*-glycosylation, followed by streptavidin-based enrichment of ConA-biotin-bound chromatin, and subsequent sequencing to identify *N*-glycosylation-bound genomic regions (Fig. 2a). To control for background from endogenous biotinylation, ConA-biotin solvent (Buffer) was used as a negative control for peak calling (Fig. 2a). Treatment with NGI-1, a specific inhibitor of the oligosaccharyltransferase (OST) complex that catalyzes *N*-glycans attachment to asparagine residues ^2,19^, significantly reduced chromatin-associated *N*-glycosylation signals, supporting the specificity of the assay (Extended Data Fig. 3a).

Analysis of GlycoChIP-seq peaks revealed that *N*-glycosylation was predominantly distributed in intergenic regions, introns, and exons (Extended Data Fig. 3c). Lamina-associated domains (LADs) are genomic regions in close contact with the nuclear lamina and INM, corresponding to chromatin at the nuclear periphery ^17,20^. By dividing the genome into LAD and non-LAD regions, we observed a marked enrichment of *N*-glycosylation within LADs (Fig. 2b, c and Extended Data Fig. 3b). To further characterize its nuclear distribution, we reconstructed the three-dimensional chromatin structure using CpG signal density as a spatial reference—given that CpG-poor regions are typically located in perinuclear and nucleolar domains (see materials and methods) ^21^. *N*-glycosylation signal intensity showed a negative correlation with CpG density, suggesting preferential localization to both perinuclear and nucleolar compartments (Fig. 2d and Extended Data Fig. 4a). This is consistent with our findings that INM proteins (Fig. 1 and Extended Data Fig. 1), as well as the nucleolar protein NOP56, are *N*-glycosylated in mESCs (Extended Data Fig. 4b).

To validate the nuclear localization of *N*-glycosylated INM proteins and investigate their potential binding to chromatin, we performed ChIP-seq for the INM proteins. TMPO (also known as Lap2β), which is known to interact with both perinuclear chromatin and the nuclear lamina, contains an *N*-glycosite within its nucleoplasmic domain ^22-25^. Building on this finding, we specifically characterized the chromatin-binding profile of its *N*-glycosylated form by performing serial ChIP (re-ChIP) in Flag-TMPO-WT-GFP mESCs (constructed as described in Fig. 3 and Extended Data Fig. 5). Chromatin was first immunoprecipitated with anti-Flag antibodies, followed by competitive elution with Flag peptides, and subsequently re-immunoprecipitated using ConA-biotin, to enrich *N*-glycosylated chromatin fragments (Fig. 2e). Both TMPO ChIP-seq and re-ChIP-seq peaks were predominantly located in intergenic regions (Extended Data Fig. 3d, e). Further analysis revealed significant enrichment of *N*-glycosylated TMPO in LADs (Fig. 2f).

**Fig. 3.**
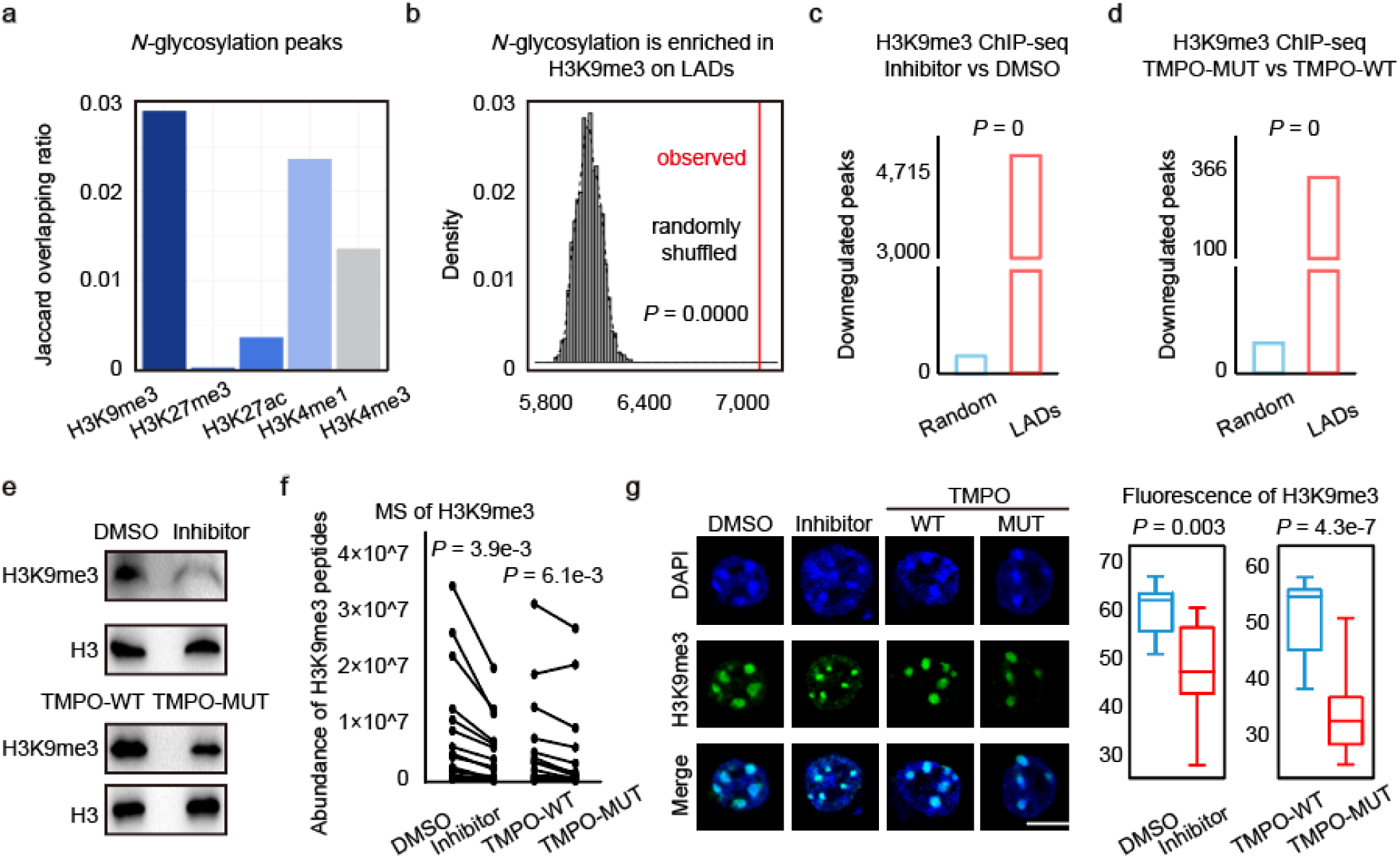
*N*-glycosylation inhibition and INM protein TMPO *N*-glycosite mutation both downregulate H3K9me3. **a**, Bar graph showing the overlapping ratios (Jaccard statistics) of *N*-glycosylated peaks with peaks of different chromatin modifications. **b**, Permutation test showing *N*-glycosylation is enriched in H3K9me3 on LADs, *P*-value by one-tailed test based on the assumption of normal distribution. **c**, Bar graph showing enrichment of downregulated H3K9me3 peaks after *N*-glycosylation inhibition in LADs, *P*-value by one-tailed test based on the assumption of normal distribution. **d**, Bar graph showing enrichment of downregulated H3K9me3 peaks after TMPO *N*-glycosite mutation in LADs, *P*-value by one-tailed test based on the assumption of normal distribution. **e**, Blotting of H3K9me3 before and after *N*-glycosylation inhibition (top), blotting of H3K9me3 before and after TMPO *N*-glycosite mutation (bottom), H3 was shown as loading control, these experiments were performed three times, with similar results. **f**, Quantitative mass spectrometry analysis of H3K9me3 peptides, *P*-values by two-tailed t-test. **g**, Immunofluorescence (left) and quantitative analysis of fluorescence signals (right) of H3K9me3, scale bar is 5 μm, *P*-values by two-tailed t-test.

In summary, immunofluorescence (Fig. 1) and established GlycoChIP-seq (Fig. 2) reveal that *N*-glycosylation localizes to the inner nuclear membrane and associates with chromatin. Meanwhile, re-ChIP experiments show that the *N*-glycosylated INM protein TMPO can also be enriched in chromatin.

## *N*-glycosylation inhibition and INM protein TMPO *N*-glycosite mutation both downregulate H3K9me3

To explore the role of *N*-glycosylation in chromatin, we analyzed its overlap with various chromatin modifications. Nearly 20% of *N*-glycosylation peaks overlap with H3K9me3, 10% with H3K4me1, 3% with H3K4me3, 2% with H3K27ac, and only 0.1% with H3K27me3. Jaccard statistics further confirmed that H3K9me3 exhibits the highest proportion of overlap with *N*-glycosylation (Fig. 3a). This observation is consistent with the established association of the inner nuclear membrane with H3K9me3-marked heterochromatin at the nuclear periphery ^18^. The observed co-enrichment with H3K4me1 may be explained by the ability of nuclear pore complexes (NPCs), which are connected to the INM, to bind enhancers at the nuclear periphery and facilitate enhancer–promoter looping for gene activation ^18^. As H3K9me3 is distributed in both LAD and non-LAD regions ^18,24^, further analysis showed that *N*-glycosylation was more prominently enriched at H3K9me3-marked regions within LADs than in non-LAD regions (Fig. 3b).

To investigate the role of *N*-glycosylation in regulating H3K9me3, two independent strategies were employed to perturb *N*-glycosylation. The first involved pharmacological inhibition using NGI-1, a small-molecule inhibitor that blocks the transfer of *N*-glycans to peptides. The second strategy entailed site-directed mutagenesis of *N*-glycosites on INM proteins. Given that TMPO interacts with both chromatin and the nuclear lamina ^22-25^, and that our data confirm its *N*-glycosylated form is associated with chromatin (Fig. 2), we selected TMPO as a representative INM protein. We first knocked down endogenous *Tmpo* in wild-type mESCs, followed by separate rescue with either wild-type TMPO (TMPO-WT) or a *N*-glycosite mutant (TMPO-MUT) (Extended Data Fig. 5a, b). Clonal lines expressing TMPO-MUT at levels comparable to TMPO-WT were selected for further analysis (Extended Data Fig. 5c). Although *N*-glycosite mutation markedly reduced *N*-glycosylated signals (Extended Data Fig. 5d, e), it did not alter the localization of TMPO (Extended Data Fig. 5f).

Four complementary experiments, including ChIP-seq, Western blotting, mass spectrometry, and immunofluorescence staining, were then employed to investigate the changes in H3K9me3 upon *N*-glycosylation inhibition and TMPO *N*-glycosite mutation. ChIP-seq results revealed a decrease in H3K9me3 signals following both treatments (Extended Data Fig. 6a, b). Further analysis revealed that H3K9me3 peaks, which were downregulated after *N*-glycosylation inhibition, were also downregulated in response to TMPO *N*-glycosite mutation (Extended Data Fig. 6c). Notably, with LAD regions remaining unaffected under the TMPO *N*-glycosite mutation and exhibiting slight changes after *N*-glycosylation inhibition, which may be attributed to the global effects of the inhibitor (Extended Data Fig. 6d, e), the downregulated H3K9me3 signals were significantly enriched in LAD regions (Fig. 3c, d). The reduced H3K9me3 signals under both conditions were further validated by Western blotting, mass spectrometry, and immunofluorescence staining (Fig. 3e-g).

Overall, multiple experimental approaches provide solid evidence that H3K9me3 is downregulated by both *N*-glycosylation inhibition and TMPO *N*-glycosite mutation.

## *N*-glycosylation inhibition and TMPO *N*-glycosite mutation both reactivate LINE-1 retrotransposons and lead to genomic instability

Given the physiological significance of H3K9me3 ^26,27^, we examined the genomic elements associated with *N*-glycosylation-regulated H3K9me3 to further investigate its biological relevance. Notably, *N*-glycosylation peaks, differential H3K9me3 peaks resulting from the two treatments, and co-localized regions of *N*-glycosylation and H3K9me3 were predominantly enriched in genomic elements such as non-LTRs and LTRs (Fig. 4a and Extended Data Fig. 6f, g). Non-LTRs, short for non-long terminal repeats, are a class of retrotransposons that include long and short interspersed elements (LINEs and SINEs, respectively), based on their size and autonomy ^28^. Among them, long interspersed nuclear element-1 (LINE-1) retrotransposons make up approximately 17% of the genome and represent the only currently autonomous retrotransposons in human ^29,30^. Their retrotransposition can result in insertional mutagenesis, chromosomal rearrangements, and genomic instability, all of which are implicated in various diseases ^31,32^. Several H3K9me3-associated proteins, including SETDB1, SUV39H1/2, and the human silencing hub (HUSH) complex, have been reported to repress LINE-1 retrotransposition ^30,31^.

**Fig. 4.**
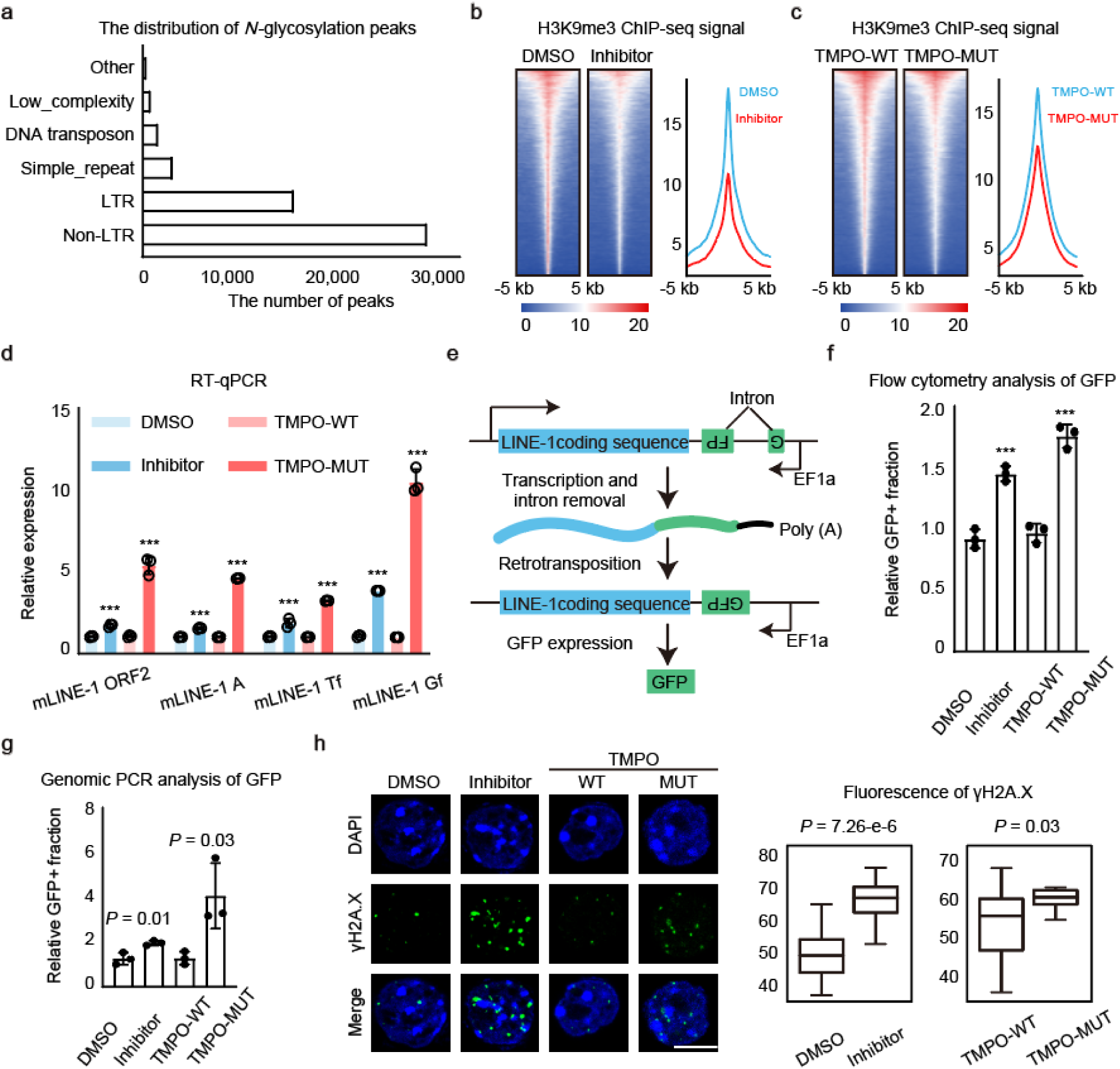
*N*-glycosylation inhibition and TMPO *N*-glycosite mutation both reactivate LINE-1 retrotransposons and lead to genomic instability. **a**, Bar graph showing the distribution of *N*-glycosylation peaks. **b**, Heat maps illustrating the density of H3K9me3 ChIP-seq reads on H3K9me3 peaks that overlap with LINE-1 before and after *N*-glycosylation inhibition in mESCs, and read-count tag density pileups of H3K9me3 profiles on H3K9me3 peaks that overlap with LINE-1. **c**, Heat maps illustrating the density of H3K9me3 ChIP-seq reads on H3K9me3 peaks that overlap with LINE-1 before and after TMPO *N*-glycosite mutation, and read-count tag density pileups of H3K9me3 profiles on H3K9me3 peaks that overlap with LINE-1. **d**, Quantitative PCR analysis of LINE-1 elements after different treatments, data were presented as mean ± SD (n = 3 biologically independent samples), *P*-values by two-tailed t-test, significance levels were indicated by asterisks: \**P* < 0.05; \*\**P* < 0.01; \*\*\**P* < 0.001 (not significant, denoted as n.s.). **e**, Schematic of the LINE-1 retrotransposition reporter assay. **f**, Bar graph showing the relative GFP+ fraction measured by flow cytometry, data were presented as mean ± SD (n = 3 biologically independent samples), *P*-values by two-tailed t-test, significance levels were indicated by asterisks: \**P* < 0.05; \*\**P* < 0.01; \*\*\**P* < 0.001 (not significant, denoted as n.s.). **g**, Bar graph showing the relative GFP+ fraction measured by genomic PCR, data were presented as mean ± SD (n = 3 biologically independent samples), *P*-values by two-tailed t-test. **h**, Immunofluorescence (left) and quantitative analysis of fluorescence signals (right) of γH2A.X, scale bar is 5 μm, *P*-values by two-tailed t-test.

To investigate the role of *N*-glycosylation-regulated H3K9me3 in LINE-1 retrotransposition, we examined H3K9me3 levels at LINE-1 regions and found that both *N*-glycosylation inhibition and TMPO *N*-glycosite mutation led to reduced H3K9me3 enrichment (Fig. 4b, c), suggesting a potential reactivation of LINE-1 elements. To validate this result, quantitative PCR with reverse transcription (RT-qPCR) using primers designed to selectively target LINE-1 subfamilies was performed, and it showed that the expression of LINE-1 was upregulated following both treatments (Fig. 4d). To functionally assess retrotransposition, we established a stable LINE-1 reporter in mESCs, in which a GFP gene interrupted by an intron and driven by the EF1α promoter was inserted in antisense orientation into the 3′-UTR of a full-length LINE-1. Successful retrotransposition leads to transcription, splicing to remove the intron, reverse transcription, and genomic integration, restoring GFP expression (Fig. 4e). Upon NGI-1 inhibitor treatment, both flow cytometry and genomic PCR showed an increased proportion of GFP-positive, indicating elevated retrotransposition activity (Fig. 4f, g). The experiment with the TMPO *N*-glycosite mutation also confirmed this result (Fig. 4f, g). Given that LINE-1 activation can cause genomic instability through the generation of double-strand DNA breaks, which in turn activate the phosphorylation of histone H2A.X (γH2A.X) ^31^, we performed immunofluorescence staining and observed an increase in γH2A.X foci (Fig. 4h), indicating that *N*-glycosylation inhibition and TMPO *N*-glycosite mutation led to genomic instability.

Collectively, these results demonstrate that *N*-glycosylation inhibition and TMPO *N*-glycosite mutation both reactivate LINE-1 retrotransposons and lead to genomic instability.

## *N*-glycosylation regulates the association of SETDB1 with the inner nuclear membrane and maintains H3K9me3

To elucidate the mechanism by which *N*-glycosylation regulates H3K9me3, immunoprecipitation of TMPO coupled with mass spectrometry analysis (IP-MS) was performed. This analysis identified a subset of H3K9 methylation-associated proteins might interact with TMPO (Fig. 5a). Among known H3K9me3 methyltransferases ^26^, only SETDB1 was detected to interact with TMPO, a finding further validated by co-immunoprecipitation (Co-IP) (Fig. 5b). Notably, the interaction between TMPO and SETDB1 was weakened after *N*-glycosylation inhibition and TMPO *N*-glycosite mutation (Fig. 5c), suggesting that this interaction was regulated by *N*-glycosylation.

**Fig. 5.**
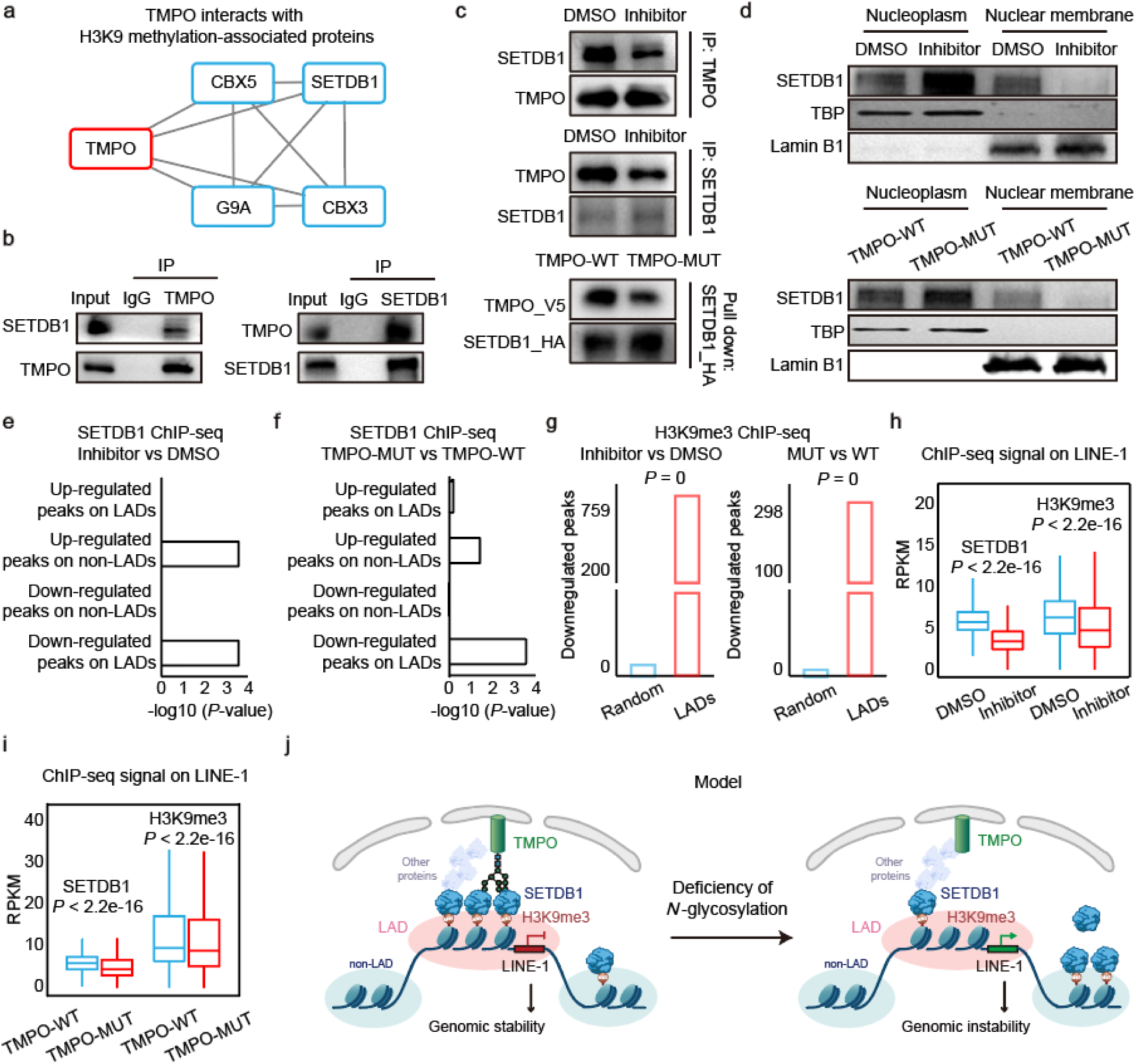
*N*-glycosylation regulates the association of SETDB1 with the inner nuclear membrane and maintains H3K9me3. **a**, Cytoscape visualization showing the interaction of TMPO with H3K9 methylation-associated proteins. **b**, Validation of the endogenous interaction between TMPO and SETDB1 by antibody-based IP in mESCs. **c**, Co-immunoprecipitation assays demonstrating that the interaction between TMPO and SETDB1 weakens after *N*-glycosylation inhibition (top and middle) and pulldown assay showing that the interaction between TMPO and SETDB1 weakens after TMPO *N*-glycosite mutation (bottom), these experiments were performed three times, with similar results. **d**, Blotting of SETDB1 from different subcellular locations, TBP and Lamin B1 were shown as loading control, these experiments were performed three times, with similar results. **e**, Bar graph showing the distribution of differential SETDB1 ChIP-seq peaks after *N*-glycosylation inhibition. **f**, Bar graph showing the distribution of differential SETDB1 ChIP-seq peaks after TMPO *N*-glycosite mutation. **g**, Bar graphs showing the enrichment of SETDB1-dependent H3K9me3 peaks downregulated after different treatments in LADs, *P*-values by one-tailed test based on the assumption of normal distribution. **h**-**i**, Box plots showing ChIP-seq signals of SETDB1 and H3K9me3 on LINE-1, *P*-values by Wilcoxon test. **j**, Model of *N*-glycosylation of INM protein in regulation of H3K9me3.

Given that INM proteins can modulate the recruitment of other proteins to the inner nuclear membrane ^25^, we further investigated whether the regulation of TMPO-SETDB1 interaction by *N*-glycosylation could influence the association of SETDB1 with the INM. Nuclear membrane and nucleoplasmic fractions were extracted before and after either *N*-glycosylation inhibition or TMPO *N*-glycosite mutation, and the results showed that SETDB1 levels decreased in the nuclear membrane fraction but increased in the nucleoplasmic fraction (Fig. 5d). To further validate the results, we divided the genome into LAD and non-LAD regions, representing genomic regions in contact with, or not in contact with, the nuclear membrane, respectively ^20^. ChIP-seq analysis showed that the decreased peaks of SETDB1 were enriched in LAD regions, whereas increased peaks were primarily located in non-LADs (Fig. 5e, f). These results demonstrate that *N*-glycosylation inhibition and TMPO *N*-glycosite mutation both result in decreased SETDB1 localization at the nuclear membrane and increased accumulation in the nucleoplasm, indicating that *N*-glycosylation regulates the association of SETDB1 with the INM.

To further explore the effect of *N*-glycosylation–regulated association of SETDB1 with the INM on H3K9me3, genome-wide H3K9me3 regions were classified into SETDB1-dependent and - independent regions. An integrated analysis revealed that both *N*-glycosylation inhibition and TMPO *N*-glycosite mutation resulted in a decrease in H3K9me3 signals in SETDB1-dependent regions (Extended Data Fig. 6h). In contrast, H3K9me3 signals remained unchanged in SETDB1-independent regions after TMPO *N*-glycosite mutation, while a slight decrease was observed after *N*-glycosylation inhibition, which may be attributed to the global regulatory effects of the inhibitor on the cell (Extended Data Fig. 6h). Moreover, the downregulated SETDB1-dependent H3K9me3 signals observed after *N*-glycosylation inhibition and TMPO *N*-glycosite mutation were also enriched in LAD regions (Fig. 5g), consistent with the LAD enrichment of downregulated H3K9me3 under the same conditions (Fig. 3c, d). Further analysis revealed that both SETDB1 and H3K9me3 signals decreased at LINE-1 elements following either treatment (Fig. 5h, i).

In conclusion, these findings uncover a regulatory role for *N*-glycosylation in the association of SETDB1 with the INM and the maintenance of H3K9me3 (Fig. 5j).

## Canonical *N*-glycan biosynthetic machinery in ER contributes to the *N*-glycosylation of inner nuclear membrane proteins

To determine the role of canonical *N*-glycan biosynthetic machinery in the *N*-glycosylation of INM proteins, key steps in *N*-glycan biosynthesis were perturbed. *N*-glycan synthesis is known to be initiated in the ER by the transfer of GlcNAc-1-P from UDP-GlcNAc to dolichol phosphate (Dol-P), forming dolichol pyrophosphate *N*-acetylglucosamine (Dol-P-P-GlcNAc), a reaction inhibited by tunicamycin (Fig. 6a) ^1^. Immunofluorescence staining of isolated nuclei following tunicamycin treatment revealed a marked reduction in ConA signals (Fig. 6b).

**Fig. 6.**
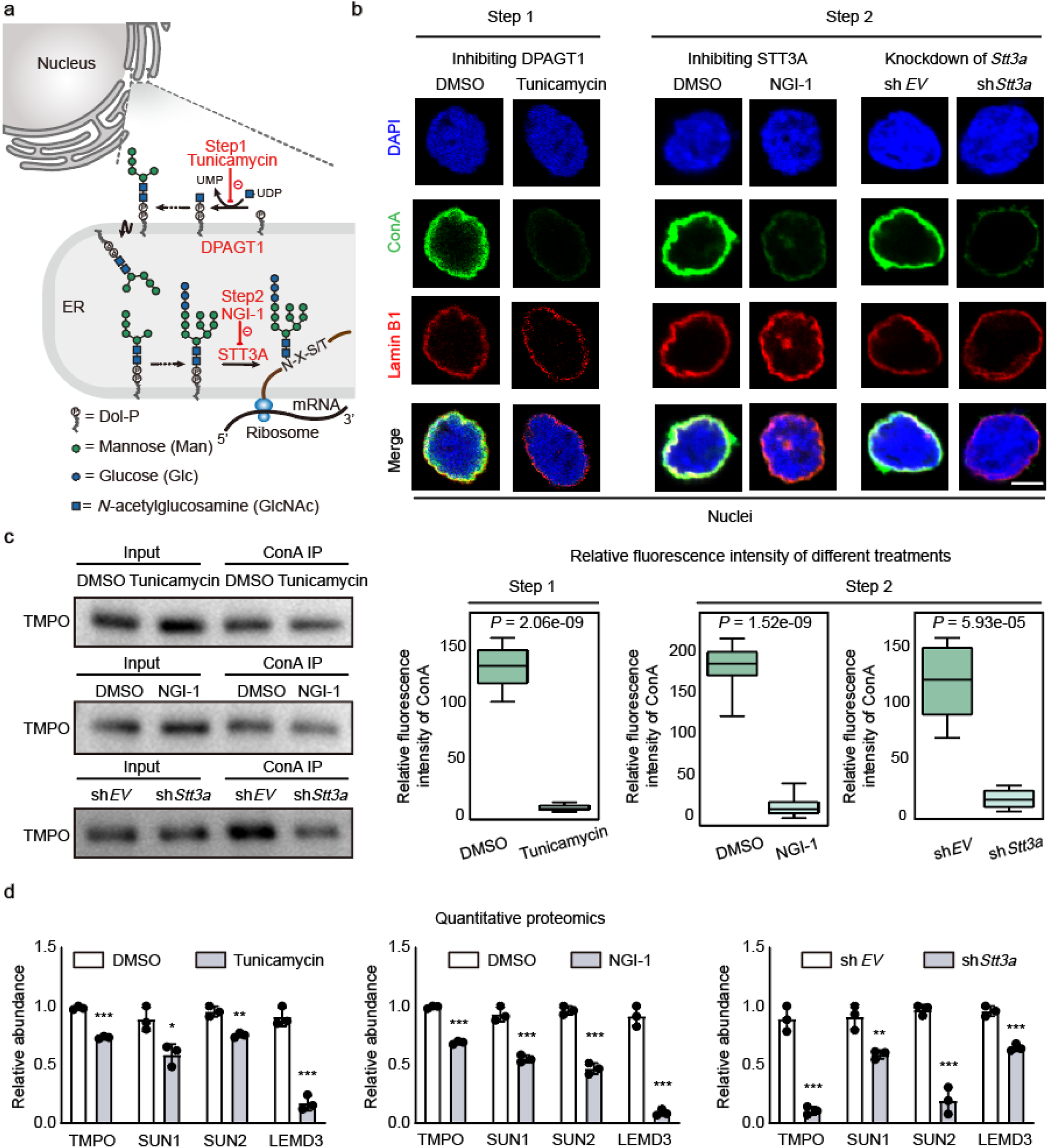
Canonical *N*-glycan biosynthetic machinery in ER contributes to the *N*-glycosylation of INM proteins. **a**, An overview of major steps in *N*-glycosylation. **b**, Immunofluorescence (top) and quantitative analysis of fluorescence signals (bottom) of ConA subjected to tunicamycin and NGI-1 treatment, as well as gene knockdown, scale bar is 5 μm, *P*-values by two-tailed t-test. **c**, Blotting analysis to detect changes in the content of the INM protein TMPO in ConA-enriched *N*-glycosylated nuclear proteome, these experiments were performed three times, with similar results. **d**, Relative abundance of INM proteins in ConA-enriched *N*-glycosylated nuclear proteome detected by quantitative proteomics, data were presented as mean ± SD (n = 3 biologically independent samples), *P*-values by two-tailed t-test, significance levels were indicated by asterisks: \**P* < 0.05; \*\**P* < 0.01; \*\*\**P* < 0.001 (not significant, denoted as n.s.).

The synthesized lipid-glycan precursor subsequently undergoes multiple processing steps before being transferred to peptides by the oligosaccharyltransferase (OST) complex, a process inhibited by NGI-1. Consistently, NGI-1 treatment also reduced ConA signals in isolated nuclei (Fig. 6b). Moreover, knockdown of *Stt3a*, a catalytic subunit of the OST complex, resulted in a decreased ConA signals (Fig. 6b and Extended Data Fig. 7a). To further validate these findings, nuclear proteins were extracted from cells subjected to these three treatments, and ConA-enriched glycoproteins were purified under denaturing conditions to eliminate protein-protein interactions (Extended Data Fig. 7b, c). Western blotting and mass spectrometry analysis of ConA-enriched *N*-glycosylated nuclear proteome revealed a reduction in the content of INM proteins (Fig. 6c, d). In contrast, inhibition of downstream *N*-glycan processing in the Golgi using kifunensine and swainsonine—specific inhibitors of α-mannosidase I and II ^2^, respectively—had no detectable effect on ConA signals (Extended Data Fig. 7d, e).

Together, these results show that the canonical *N*-glycan biosynthetic machinery in the ER contributes to the *N*-glycosylation of INM proteins.

## Discussion

While polysaccharides are known to play essential roles at the cell membrane and in the extracellular space, their presence and function in the nucleus have not previously been reported. In this study, we demonstrate that, with the support of the canonical biosynthetic machinery in the ER, *N*-glycans—a class of polysaccharides—can modify inner nuclear membrane proteins and exist in the nucleus. *N*-glycosylated INM proteins play an important role in the maintenance of H3K9me3-heterochromatin and genomic stability. These findings establish a link between polysaccharides and chromatin regulation, broadening their biological roles and advancing our understanding of nuclear regulatory mechanisms.

### Potential mechanisms for the targeting of *N*-glycosylation to the nucleus

Among currently recognized saccharides, *N*-acetylglucosamine (GlcNAc) is uniquely capable of modifying nuclear proteins as a single monosaccharide, owing to the nuclear localization of its transferase enzyme, O-GlcNAc transferase (OGT). In contrast, enzymes involved in *N*-glycosylation have been identified in the ER and Golgi apparatus. The canonical *N*-glycosylation pathway operates as a secretory process initiated in the ER, where a lipid-glycan precursor— comprising three glucose (Glc), nine mannose (Man), and two *N*-acetylglucosamine (GlcNAc) residues attached to dolichol phosphate—is transferred to polypeptides by the OST complex (Fig. 6a). Following this transfer, glucose residues are sequentially removed by two α-glucosidases (α-Glc I–II), and one mannose residue is removed by the ER α-mannosidase (ER α-Man). After passing a quality-control checkpoint, the glycoprotein is transported to the Golgi apparatus for further maturation ^9^. The majority of glycoproteins exiting the ER carry *N*-glycans containing either eight or nine mannose residues (Man8GlcNAc2 or Man9GlcNAc2) ^1^.

Given that (i) the *N*-glycans identified on inner nuclear membrane proteins are Man8GlcNAc2 or Man9GlcNAc2 (Fig. 1b), (ii) the results in Fig. 6 demonstrate that the canonical *N*-glycan biosynthetic machinery in the ER contributes to the *N*-glycosylation of INM proteins, and (iii) newly synthesized INM proteins are initially inserted into the ER, subsequently transported to the nucleus and ultimately targeted to the INM ^33^, we speculate that *N*-glycans are added to INM proteins in the ER and then co-translocated with these proteins into the nucleus during INM targeting, suggesting that the presence of *N*-glycans within the nucleus is likely associated with the INM targeting of INM proteins. The current model for INM targeting involves multiple steps: newly synthesized INM proteins are first integrated into the ER membrane, diffuse through the continuous membrane system comprising the ER and the outer nuclear membrane (ONM), translocate through the peripheral or central channels of nuclear pore, and are retained in the INM via interactions with the nuclear lamina or chromatin (Extended Data Fig. 8a) ^34-38^. To preliminarily investigate the mechanism of *N*-glycosylation targeting to the nucleus, we disrupted certain steps in the targeting process of INM proteins to INM. Previous studies have established that Atlastins, membrane-bound GTPases of the ER, play an essential role in the diffusion and subsequent targeting of proteins to the inner nuclear membrane. Among them, ATL2 has been identified as a key regulator of this process ^35^. We knocked down *Atl2*, and results suggest that the diffusion and the INM targeting of *N*-glycosylated INM proteins likely also require Atlastin (Extended Data Fig. 8). Furthermore, studies have found that when INM proteins translocate through the nuclear pores, proteins such as NUP35, NUP93, NDC1, and NUP210 influence INM targeting ^33,34^. We knocked down these genes, and results suggest that these nuclear pore complexes are also likely to be involved in the targeting of *N*-glycosylated INM proteins to the INM (Extended Data Fig. 8). These results suggest that *N*-glycosylation is targeted to the nucleus, potentially linked to the targeting process of INM proteins to the INM.

However, the known catalytic subunit of OST, STT3A, is localized within the lumen of the endoplasmic reticulum (Fig. 6a) ^1,5^, whereas the *N*-glycosite of TMPO lies within its nucleoplasmic domain ^22-25^, and we demonstrate that the *N*-glycosylated form of TMPO is enriched on chromatin within the nucleoplasm (Fig. 2). This raises the question of whether TMPO is initially *N*-glycosylated within the ER lumen and subsequently flipped to the cytoplasmic face of the ER membrane via an uncharacterized mechanism, or if *N*-glycosylation occurs directly at the cytoplasmic face through an alternative, yet-to-be-identified pathway. In addition to these scenarios, other possibilities may also exist, and the precise molecular mechanisms underlying these processes remain to be elucidated.

### The roles of *N*-glycosylation in the nucleus

Abnormalities in INM proteins have been implicated in a range of disorders collectively referred to as “nuclear envelopathies” ^39^. Given our finding that *N*-glycans can modify INM proteins across diverse cell types (Fig. 1), as well as our observation that deficiency in *N*-glycosylation leads to genomic instability (Fig. 4), an interesting future direction would be to investigate whether the above mechanisms are involved in disease progression.

Over half of the vertebrate genome is packaged into transcriptionally repressed H3K9me3 heterochromatin, which is important for silencing transposons and preventing aberrant transcription. While proteins such as TNRC18 and ZNF512 ^40,41^, as well as RNA ^42^, have been shown to regulate H3K9me3, our study reveals that polysaccharides—previously identified only outside the nucleus—also function within the nucleus to modulate H3K9me3. Importantly, these polysaccharides specifically maintain H3K9me3-marked heterochromatin in LADs (Fig. 3), a genomic region essential for the silencing of LINE-1 retrotransposons and the maintenance of genomic stability ^17^. Moreover, *N*-glycans modify inner nuclear membrane proteins across multiple species and cell types (Fig. 1), suggesting a cell-type-invariant and conserved mechanism by which *N*-glycosylation contributes to the maintenance of heterochromatin in LAD regions. In addition to non-LTR regions such as LINE-1, we also observed overlapping signals of *N*-glycosylation and H3K9me3 in LTR regions (Extended Data Fig. 6). Given the importance of H3K9me3-mediated repression of LTR retrotransposons in maintaining cell identity ^43,44^, it would be interesting to explore the role of *N*-glycosylation in this process.

In conclusion, our findings demonstrate that polysaccharides contribute to the maintenance of H3K9me3-heterochromatin and genomic stability within the nucleus, extending their biological significance beyond the cell membrane and extracellular matrix.

## Methods

### Antibodies

Antibodies used in this study were anti-TMPO (Proteintech, #14651-1-AP), anti-GFP (Proteintech, #50430-2-AP), anti-SETDB1 (Proteintech, #11231-1-AP), anti-H3K9me3 for ChIP-seq (Abcam, #ab8898), anti-H3K9m3 for IF (ABclonal, #A22295), anti-Lamin B1 (ABclonal, #A16909), Normal rabbit IgG (Millipore, #12-370), and anti-Phospho-Histone H2A.X (Cell Signaling Technology, #9718S). TrueBlot Anti-Rabbit IgG HRP (Rockland, #18-8816-33), TrueBlot Anti-Mouse IgG HRP (Rockland, #18-8817-33), Goat anti-rabbit-IgG (H+L)-HRP (Cell Signaling Technology, #7074), Goat anti-mouse-IgG (H+L)-HRP (Cell Signaling Technology, #7076), Goat anti-Mouse IgG (H+L) Highly Cross-Adsorbed Secondary Antibody, Alexa Fluor™ 488 (Thermo Fisher, #A-11029), Goat anti-Rabbit IgG (H+L) Highly Cross-Adsorbed Secondary Antibody, Alexa Fluor™ 594 (Thermo Fisher, #A-11037) were used as secondary antibodies.

### Cell lines

Mouse and human embryonic stem cells (ESCs), mouse pre-iPS cells, mouse embryonic fibroblasts (MEFs), and mouse totipotent-like stem cells (TLSCs) were used for the identification of nuclear *N*-glycosylation. Mouse embryonic stem cells (ESCs) were used for further downstream experiments. HEK293T cells were used for lentivirus packaging.

### Cell culture

The mouse ESC line R1 and the pre-iPSC line were cultured on gelatin-coated dishes with ESC medium ^45^. The ESC medium composition included DMEM (Hyclone, #SH30022.01), 15% (v/v) fetal bovine serum (FBS, Lonsera, #S712-012S), 0.1 mM β-mercaptoethanol (Sigma, #M6250), 2 mM L-glutamine (Thermo Fisher, #35050061), 0.1 mM nonessential amino acids (Thermo Fisher, #11140050), 1% (v/v) nucleoside mix, and LIF. Human ESC line H9 was maintained in mTeSR™ media (StemCell Technologies, #85850) on tissue culture plates coated with Matrigel (BD Bioscience, #356230). MEF cells were cultured in a medium consisting of 10% FBS and 90% DMEM medium. Mouse TLSCs were generously provided by Professor Jichang Wang at Sun Yat-sen university and cultured in the TLSC medium (TLSCM) comprising KnockOut medium, 20% KnockOut serum replacement (Thermo Fisher, #10828028), 1 x nonessential amino acids solution (Thermo Fisher, #11140050), 2 mM L-glutamine (Thermo Fisher, #25030081), 50 μg/ml BSA (Thermo Fisher, #15260037), 100 μM 2-mercaptoethanol (Thermo Fisher, #21985023), 10 ng/ml IL6 (PeproTech, #AF-200-06), 10 ng/ml sIL-6R (PeproTech, #200-06RC), 100 μg/ml L-ascorbic acid (Sigma, #A4544), 2 μM SGC0946 (Cayman Chemical, #13967), 2 μM A366 (Tocris, #5163), 3 μM AS8351 (Tocris, #6044), and primocin (InvivoGen, #Ant-pm-2) ^9^.

### Construction of knockdown cell lines

Short hairpin RNAs (shRNAs) were designed and synthesized. Then DNA sequences were inserted into the lentiviral plasmid pLKO.1. Plasmids psPAX2, pMD2.G and pLKO.1-shRNA were transiently transfected into HEK293T cells to prepare the lentiviruses using PEI (Yeasen, #40816ES02). The lentivirus supernatant was harvested. For virus infection, mESCs were seeded onto 6-well plates the day before infection. The following day, the spent medium was aspirated and replaced with 1 ml of mESC medium and 1 ml of shRNA lentivirus containing 4 µg/ml polybrene (Beyotime, #C0351). After 24 h of infection, the medium was replaced with fresh medium, and infected cells were selected with specific resistance.

### Isolation of nuclei

Gagnon et al. (2014) described an optimized protocol that cleanly recovers mammalian nuclei after processing without significant residual ER membrane attached ^46^. This method was also used by Ryan A. Flynn et al. (2021) to identify the localization of glycoRNA ^2^. We followed this protocol step by step for nuclear isolation without making any modifications.

### Protein digestion

The nuclei samples were denatured individually in 8 M urea (Sigma, #U5128)/1 M NH_4_HCO_3_ (Sigma, #11213) and subjected to ultrasonication on ice using an Ultrasonic Cell Distribution System. Subsequently, the denatured samples were subjected to centrifugation at 15,000 g for 20 min, after which the supernatants were collected for the purpose of measuring protein concentration using the BCA reagent (Beyotime, #P0012). To reduce disulfide bonds between proteins, 5 mM DTT (Sigma, #43819) solution was added to the protein solution and shaken at 160 rpm and 37 °C for 1 h. The protein was then alkylated with 15 mM iodoacetamide (Sigma, #I6125) for 30 min at room temperature in the dark, followed by the addition of 2.5 mM DTT for 10 min at room temperature. The protein samples were diluted twofold with deionized water and digested with sequencing grade trypsin (protein: enzyme, 100:1, w/w; Promega, #V5113) at 37 °C for 2 h with gentle shaking, representing the initial digestion stage. Following this, the samples were diluted fourfold with deionized water, and sequencing grade trypsin (protein: enzyme, 100:1, w/w; Promega, #V5113) was employed to digest the proteins into peptides once more through incubation at 37 °C with gentle shaking overnight, representing the second digestion. The samples were acidified with trifluoroacetic acid (TFA) (Sigma, #302031) and subjected to centrifugation at 15,000 g for 15 min in order to remove any particulate matter. The digested peptides were desalted with a C18 column and eluted with 50% acetonitrile (ACN) (Thermo Fisher Scientific, #A9554)/0.1% TFA. The peptide concentration was determined by measuring the UV absorbance at 215 nm using a DS-11 spectrophotometer (DeNovix, USA).

### Enrichment of *N*-linked intact glycopeptides

Intact glycopeptides were enriched using a mixed anion exchange column (MAX) (Waters, #186000366). Briefly, the peptides eluted from the C18 column (Waters, #WAT054955) were mixed with 100% ACN and TFA to a final concentration of 95% ACN/1% TFA. The MAX columns were activated by the sequential addition of 1 ml of 100% ACN, 100 mM triethylammonium acetate buffer (Sigma, #69372), water, and 95% ACN/1% TFA solution. This was followed by the loading and washing of the samples with a solution of 95% ACN/1% TFA. The glycopeptides were then eluted in 400 μl of 50% ACN/0.1% TFA solution, dried by vacuum concentration with an RVC 2-18 CD plus concentrator (Christ, Germany), and resuspended in 20 μl of 0.1% TFA for liquid chromatography tandem MS (LC−MS/MS) analysis.

### LC-MS/MS analysis

The detailed LC-MS/MS method was described in our previous publication ^47^. Each sample was subjected to an LC-MS/MS analysis on an Orbitrap Fusion Lumos Mass Spectrometer coupled with an Easy-nLC 1200 system (Thermo Fisher Scientific, Germany). About 1 μg intact glycopeptide was separated by an Easy-nLCTM 1200 system (Thermo Fisher Scientific, USA) with the use of a 75 μm × 50 cm Acclaim PepMap-100 C18 analytical column (Thermo Fisher Scientific, #164570) protected by a 75 μm × 2 cm trapping column (Thermo Fisher Scientific, #164946). The flow rate was maintained at 200 nl/min with mobile phase consisting of 0.1% TFA in water (A) and 0.1% TFA in 80% ACN (B). The gradient profile (240 min) for glycoproteomics was set as follows: 3–40% B for 203 min, 40–68% B for 20 min, 68–99% B for 4 min, finally, 99% B for 13 min.

The settings for intact glycopeptide analysis were set as follows: the spray voltage was set at 2.4 kV. Orbitrap MS1 spectra (AGC 4.0 × 105) were collected from 375-1800 m/z at a resolution of 120 K. Each selected MS1 peak (isolation width of 2 m/z) was fragmented by data-dependent HCD with collision energy of 20% and 33% at a resolution of 30 K (two MS2 spectra per glycopeptide). High HCD energy of 33% was used for identification of the peptide sequence, while low HCD energy of 20% was used for glycan structure analysis of the intact glycopeptide. Charge states from 2 to 7 were selected for MS/MS acquisition. A dynamic exclusion time of 10 s was used to discriminate against previously selected ions.

### Intact glycopeptide (IGP) identification and quantification

The identification of IGPs was performed by our developed software, StrucGP ^8^. IGPs analyses were performed by StrucGP using the built in glycan branch structure database from StrucGP and the mouse protein databases (Uniprot, UP000000589). The protein enzymatic digestion was set as trypsin with a maximum of two missed cleavage sites, and the potential glycosite containing peptides were screened with the N-X-S/T motif (X is any amino acid except Proline). The carbamidomethylation (C, +57.0215 Da) was as a fixed modification, and oxidization (M, +15.9949 Da) as a dynamic modification. The mass tolerances for MS1 and MS2 were set at 10 ppm and 20 ppm, respectively. The identification results were filtered with 1% False discovery rate (FDR) for both peptide sequences and glycan structures, which was estimated by the decoy peptide method and decoy spectra method, respectively. The search results were ranked based on their scores, and the peptide/glycan with the highest score was considered to be the corrected identification.

### Protein purification

A total of 50 million cells were harvested and lysed in lysis buffer (50 mM HEPES pH = 7.6, 250 mM NaCl, 0.1% NP-40, 0.2 mM EDTA, 0.2 mM PMSF, 1 x protease inhibitor cocktail) on ice. The lysates were cleared by centrifugation and the supernatant was protein extraction. The protein extraction was denatured by boiling for 10 min to disrupt protein-protein interactions. For GFP-based purification, GFP antibody was conjugated with Protein G beads (Beyotime, #P2106) by incubating in IP DNP buffer (20 mM HEPES pH = 7.6, 0.2 mM EDTA, 1.5 mM MgCl_2_, 100 mM KCl, 20% glycerol, 0.02% NP-40, 1 x protease inhibitor cocktail) overnight at 4 °C and washed twice. Then, protein extraction was added to the GFP-conjugated Protein G beads and rotated at 4 °C. After 12 h, the supernatant was removed and the GFP-conjugated Protein G beads were washed twice with IP DNP buffer. 2 x SDS loading buffer was added to the GFP-conjugated Protein G beads and the protein was eluted by boiling 5 min at 95 °C. SDS-PAGE analysis was applied to assess purification efficiency and the remaining protein was kept in -80 °C. For Flag purification, anti-Flag magnetic beads (Beyotime, #P2115) were added to the protein extraction and incubated overnight at 4 °C. The next day, the beads were washed twice with IP DNP buffer, and the supernatant was removed. 3 x Flag peptide (Beyotime, #P980) was added for competitive elution. After three rounds of elution, the eluate was subjected to ultrafiltration to remove the 3 x Flag peptide. The purified protein was then frozen at -80 °C for subsequent PNGase F treatment.

### ConA-enriched *N*-glycosylated proteins

25 million cells were used for isolation of nuclei, and the extracted protein was denatured by boiling for 10 min at 95 °C. Denatured protein was incubated overnight at 4 °C with agarose-bound Con A resin (Vector Laboratories, #AL-1003) and the agarose was then washed three times. 2 x SDS loading buffer was added and the protein was eluted by boiling 5 min at 95 °C. Agarose without Con A conjugated was used as control.

### The treatment of PNGase F

The treatment of PNGase F (NEB, #P0705S) was performed according to the manufacturer’s instructions. For the digestion of glycans from protein, 1 µl of PNGase F was used per 5 µg of purified protein. For isolated nuclear staining, 3 µl of PNGase F was employed.

### The treatment of inhibitors

Working stocks of inhibitors were all made in DMSO at the following concentrations: 118.35 nM tunicamycin (Abcam, #ab120296), 10 mM NGI-1 (Sigma, #SML1620), 10 mM Kifunensine (Sigma, #K1140), 10 mM Swainsonine (Sigma, #S9263).

### Western blot

Samples underwent electrophoresis on SDS-PAGE gels and were subsequently transferred to PVDF membranes (BIO-RAD, #1620177). Following this, the membranes were blocked with 5% BSA at room temperature. After a one-hour incubation period, the membranes were incubated with primary antibodies overnight at 4 °C, followed by three washes with TBST and subsequent incubation with peroxidase-labeled secondary antibodies for 1 h at room temperature. Following another three washes with TBST, bands were visualized using an ECL substrate (BIO-RAD, #1705061) and imaged with a CCD camera.

### Lectin blot

All procedures closely resemble those of western blot, except that in lectin blot, the primary antibody was lectin conjugated with biotin (Vector, #B-1005-5) and the secondary antibody was peroxidase-labeled streptavidin (Vector, #A-2014).

### Immunofluorescence and image analysis

Cells were cultured on gelatin-coated glass and subsequently fixed in 4% paraformaldehyde (PFA, Solarbio, #P1110) for 15 min. After washing with PBS, cells were permeabilized with 0.25% Triton X-100 (AMERSCO, #0694-1L) and blocked with 10% bovine serum albumin. After 1 h, cells were incubated with primary antibodies in 3% BSA overnight at 2–8 °C. After washing 3 times with PBS, cells were incubated with secondary antibodies for 1 h. After two washes of PBS, cells were stained with DAPI for 10 min at room temperature. Confocal microscopy was utilized to image the fixed cells. 3D reconstructions were performed using IMARIS software (Bitplane AG, Switzerland). For immunolocalization in isolated nuclei, staining nuclei devoid of the surrounding cytoplasm followed the methods developed by Nagaraj et al., 2017 ^48^. Specifically, cells were collected and washed with PBS, transferred with minimal amounts of PBS to a microdrop of 0.01 nM HCl/0.1% Tween 20 (Amresco, #0777-1L) in distilled sterile water on a glass slide. After 2 min, the cells were examined under a microscope, carefully removed the cytoplasm. The slides were washed with several rounds of PBS to remove the remaining cytoplasm. The attached nuclei were fixed with 4% paraformaldehyde. The following steps were the same as immunofluorescence. The signal intensity of immunofluorescence was quantitated by ImageJ (https://imagej.net/Fiji).

### ChIP-seq

Cells were crosslinked with 1% formaldehyde (Sigma, #F8775) for 10 min at room temperature and then quenched with 125 mM glycine (Sigma, #G7126) for 5 min. After then, the cells were collected and incubated in lysis buffer I (50 mM HEPES-KOH pH = 7.5, 140 mM NaCl, 1 mM EDTA, 10% glycerol, 0.5% NP-40, 0.25% Triton X-100, protease inhibitors). After 10 min, the cells were collected, resuspended in lysis buffer II (10 mM Tris-HCl pH = 8.0, 200 mM NaCl, 1 mM EDTA, 0.5 mM EGTA, protease inhibitors) and rotated for 10 min. For sonication, the cells were collected, and resuspended in sonication buffer (20 mM Tris-HCl pH = 8.0, 150 mM NaCl, 2 mM EDTA pH = 8.0, 0.1% SDS, and 1% Triton X-100, protease inhibitors). Sonicated lysates were cleared once by centrifugation at 16,000 g for 10 min at 4 °C and the supernatant was transferred to 15 ml conical tube. 50 μl supernatant was saved as input. The remainder of the mixture was incubated with magnetic beads bound with antibody to enrich for DNA fragments overnight at 4 °C. The next day, beads were washed with wash buffer (50 mM HEPES-KOH pH = 7.5, 500 mM LiCl, 1mM EDTA pH = 8.0, 0.7% Na-Deoxycholate, 1% NP-40) and followed with TE buffer (10 mM Tris-HCl pH = 8.0, 1 mM EDTA, 50 mM NaCl). Beads were removed by incubation at 65 °C for 30 min in elution buffer (50 mM Tris-HCl pH = 8.0, 10 mM EDTA, 1% SDS) and supernatant was reverse crosslinked overnight at 65 °C. To purify eluted DNA, 200 μl TE was added to dilute SDS and 8 μl 10 mg/ml RNase A (Thermo Fisher, #EN0531) was added to degrade RNA. After 2 h, protein was degraded by addition of 4 μl 20 mg/ml proteinase K (Thermo Fisher, #25530049) and incubation at 55 °C for 2 h. Phenol: chloroform: isoamyl alcohol extraction (G-CLONE, #EX0128) was performed followed by an ethanol precipitation. The DNA pellet was then resuspended in 50 μl TE. Library was performed with NEBNext Ultra II DNA library kit (NEB, #E7645). Two biological replicates were performed for each cell line.

### GlycoChIP-seq

The crosslinking, lysis, and sonication processes of GlycoChIP-seq were the same as those for ChIP-seq. For the experimental group, chromatin was incubated with ConA-biotin (Vector, #B-1005-5) on a rotator at 4 °C overnight to enrich *N*-glycosylation. For the control group, the ConA-biotin solvent (buffer) was used to exclude the effects of endogenous biotinylation. Dynabeads MyOne Streptavidin T1 (Invitrogen, #65602) were added to enrich biotin. The remaining steps, including elution, reverse cross-linking, DNA purification, and library preparation were similar to those used in ChIP-seq. Two biological replicates were performed for each cell line.

### RNA isolation and RT-qPCR

Cell pellets were homogenized in RNAzol reagent (MRC, #RN190-500) and processed according to the manufacturer’s instructions. For RT-qPCR, RT SuperMix (Vazyme, #R122-01) was used for reverse transcription. All qPCR reactions were performed as 12 µl reactions using 2 x chamQ mix (Vazyme, #Q711-03). Relative abundance was determined using 2−ΔΔCt. Three biological replicates were conducted for each experiment.

### Co-immunoprecipitation and IP-MS

10 million cells were collected and lysed with RIPA buffer, followed by centrifugation and collection of the supernatant. Endogenous TMPO was immunoprecipitated with 5 µg of TMPO antibody pre-bound to Protein G agarose and co-immunoprecipitated SETDB1 was identified by western blot with the antibody of SETDB1. The immunoprecipitation of SETDB1 was performed in the same manner, except that five times the amount of cells were used. IP-MS was performed following the method described in the literature ^49^, using 25 µg of TMPO antibody, with IgG as a control. Two biological replicates were conducted.

### Extraction of histone

Histone extraction was performed based on the method described in the literature, with appropriate modifications ^50^. 10^7^ cells were resuspended in extraction buffer (10 mM HEPES pH = 7.9, 10 mM KCl, 1.5 mM MgCl_2_, 0.34 M sucrose, 10% glycerol, and 1 x protease and phosphatase inhibitors) with 0.2% NP-40 and incubated on ice for 10 min. The supernatant was removed by centrifugation, and the pellet (the nuclei) was washed with extraction buffer (without NP-40) and incubated on ice for 1 min. After centrifugation, the supernatant was completely removed. The nuclei pellet was lysed in no-salt buffer (3 mM EDTA, 0.2 mM EGTA) and vortexed intermittently for 1 min. The mixture was then incubated on a rotator at 4 °C for 30 min. The supernatant, containing the nucleoplasm, was removed, and the pellet, containing the chromatin, was retained. The chromatin pellet was re-suspended in High-Salt Solubilization buffer (50 mM Tris pH = 8.0, 2.5 M NaCl, 0.05% NP-40) and vortexed for 2 min. The mixture was incubated on a rotator at 4 °C for 30 min, followed by centrifugation. The supernatant containing extracted histones was dialyzed and then subjected to mass spectrometry analysis.

### Extraction of nuclear membrane and nucleoplasmic fraction

The extraction was performed following the method described in the literature with some modifications ^20^. Cells were resuspended in hypotonic lysis buffer (10 mM HEPES pH = 7.9, 1.5 mM MgCl₂, 10 mM KCl) and subjected to Dounce homogenization to isolate nuclei. Isolated nuclei were resuspended in Nuclear Envelope Buffer 1 (10 mM Tris pH = 8.5, 0.1 mM MgCl₂, 5 mM β-ME, 10% sucrose, DNase I) and incubated at room temperature for 10 min. After centrifugation, the supernatant was collected as the nucleoplasmic fraction, while the pelleted nuclear membrane was resuspended in RIPA buffer.

### LINE-1 GFP retrotransposition assay

The LINE-1-GFP retrotransposition reporter was kindly provided by Professor Nian Liu, and the retrotransposition efficiency was analyzed according to a previously reported protocol with some modifications ^25^. Briefly, cells were plated in 6-well plates and transfected with the LINE-1 GFP retrotransposition reporter vector. After selection with BSD for 7 days, the cells were passaged and treated with either DMSO or NGI-1. GFP expression was assessed by flow cytometry and genomic PCR. Genomic DNA was extracted following the manufacturer’s instructions (TIANGEN, #DP304-03). A total of 300 ng of genomic DNA per sample was digested with 50 units of PspGI (NEB, #R0611S) at 75 °C to specifically cleave the intron within the GFP cassette. The reaction mixture was then used for qPCR with primers flanking the intron region of the GFP cassette. Since PspGI digestion prevents amplification of the unspliced LINE-1 GFP reporter, only newly integrated GFP cassettes, in which the intron was removed during the retrotransposition process, can be amplified by PCR.

### ChIP-seq data analysis

High-confidence reads of ChIP-seq data were obtained using fastp (v0.20.0) with default parameters and mapped to the mouse genome mm9 using Bowtie2 (v2.5.1) with parameters ’--sensitive -p 24’ ^51,52^. PCR duplicated fragments were filtered using Picard (v2.22.8) (http://broadinstitute.github.io/picard). Unmapped and multi-mapped reads were filtered out and reads mapping to chrM were removed. Unique reads were filtered by MAPQ > 30. We used deepTools (v3.5.0) bamCoverage to generate bigWig files with RPKM normalization ^53^. SAM files were converted to BAM format for peak calling using SAMtools (v1.9). Peaks were identified using the MACS2 (v2.2.4) callpeak pipeline ^54^. For GlycoChIP-seq, peak calling was performed using ConA-biotin solvent (Buffer) as the control.

Enriched peak regions generated by MACS2 (v2.2.4) were used as input for edgeR (v4.2.1) to identify differential peaks from ChIP-seq data ^55^. Peaks were annotated using the Bioconductor package ChIPSeeker (v1.26.0). For mapping peaks to gene features, we analyzed the distribution of peaks across the genome for each ChIP-seq dataset. Heatmaps and profiles were generated for regions ± 5 kb from the center of peaks using deepTools (v3.5.0) plotHeatmap, and average scores for profiles were plotted to generate averaged read density around peaks using deepTools (v3.5.0) plotProfile. The Jaccard statistic, representing the ratio of the intersection of two sets to the union of the two sets, was calculated using Bedtools (v2.29.2) ^56^. The LAD regions were identified in the study by Daan Peric-Hupkes et al., (2010) (*15*).

### Spatial distribution of ConA

We reprocessed the single-cell Hi-C data of mESCs from PMID: 28682332 (GSE94489) ^57^. The samples were demultiplexed by fastq-multx (v1.3.1, https://github.com/brwnj/fastq-multx) and mapped to mm9 using dip-c (https://github.com/tanlongzhi/dip-c). During the process, the two haplotypes of each chromosome were distinguished by extracting 129S1/SvImJ and CAST/EiJ from dbSNP VCF file. Subsequently, 3D reconstruction was performed with hickit (https://github.com/lh3/hickit, ‘hickit -i cell_impute.txt -u -s 517 -Sr1m -c1 -r10m -c2 -b4m -b1m -b200k -D5 -b50k -D5 -b20k -O cell.3dg’). For quality control of the reconstructed 3D genome structures, 3 replicates of each cell were generated with random seeds. The cells with a median RMSD value < 1.5 among the 3 replicates were used for downstream analysis. As a result, G1 cells without large chromosomal aberrations were chosen. The CpG frequencies for visualizing chromatin compartments were downloaded from https://github.com/tanlongzhi/dip-c/blob/master/color/mm10.cpg.20k.txt. The visualization of the spatial distribution of CpG frequencies and ConA was performed by PyMOL (v3.0.3, https://pymol.org/).

## Data availability

Previously published raw reads were downloaded from GSE90895 (H3K9me3 ChIP-seq, H3K27me3 ChIP-seq, H3K27ac ChIP-seq, H3K4me1 ChIP-seq, H3K4me3 ChIP-seq) ^10^, GSE17051 (LADs) ^15^, PRJNA544540 (SETDB1 ChIP-seq, SETDB1-cKO DMSO H3K9me3 ChIP-seq, SETDB1-cKO 4OHT H3K9me3 ChIP-seq) ^58^, GSE94489 (Single-cell Hi-C data of mESCs) ^57^. The GlycoChIP-seq and ChIP-seq data generated during this study are available at GEO: GSE273948. This paper does not report original code. All materials are available upon contacting the corresponding author.

## Acknowledgments

We are grateful to the members of the Ding laboratory for their valuable discussions, technical advice, and support. In particular, we thank the glycosylation group for assistance with vector construction and figure optimization, Jiajun Liu for illustrating schematic diagrams, and Zibing Huang for contributing to the reconstruction of the nuclear distribution of ConA. We thank Professor Jichang Wang from Sun Yat-sen University for providing TLSCs cells, Professor Jiekai Chen from the Chinese Academy of Sciences for his advice on SETDB1 ChIP-seq and Professor Nian Liu from Tsinghua University for providing the LINE-1 retrotransposition reporter. The following funding supported this study: National Key Research and Development Program of China 2024YFA1106900 and 2023YFA1800900 (J.J.D.), National Science Foundation for Distinguished Young Scholars of China 32425022 (J.J.D.), National Natural Science Foundation of China 32170798 and 32430031 (J.J.D.), National Natural Science Foundation of China 22374117 (S.S.S), National Key Research and Development Program of China 2019YFA0905200 (S.S.S), Natural Science Foundation of Shaanxi Province 2024JC-TBZC-06 and 2023-JC-ZD-46 (S.S.S), Shaanxi Fundamental Science Research Project for Chemistry and Biology 23JHZ006 (S.S.S), Fundamental Research Funds for the Central Universities, Sun Yat-sen University 24qnpy251(X.X.T.), National Natural Science Foundation of China 32400507 (X.X.T.), China Postdoctoral Science Foundation 2024M753727 (X.X.T.), China Postdoctoral Science Foundation 2023M744082 (X.Y.L.), National Natural Science Foundation of China 32400659 (X.Y.L.), The Science and Technology Development Fund, Macau SAR 0099/2022/AFJ, and 0061/2022/A (W.C.), National Natural Science Foundation of China T2450074 (Y.M.).

## Author contributions

X.X.T. and J.J.D. designed the study; X.X.T. wrote the manuscript with input from all other authors; X.X.T., L.Q., L.Z.L., H.C.L., W.D., Y.Q.H. and Z.D.Z. conducted experiments. R.R.D., X.X.T. and X.Y.L performed the bioinformatics analysis. Y.M., W.C. participated in project guidance and provided valuable suggestions. J.J.D. and S.S.S. supervised the project.

## Competing interests

The authors declare no competing interests.

**Correspondence and requests for materials** should be addressed to Junjun Ding or Shisheng Sun.

## Supplemental Information includes three tables

Supplementary Table 1. shRNAs used in this study.

Supplementary Table 2. Primers used in this study.

Supplementary Table 3. ChIP-seq statistics.

---

**Extended Data Fig. 1.**
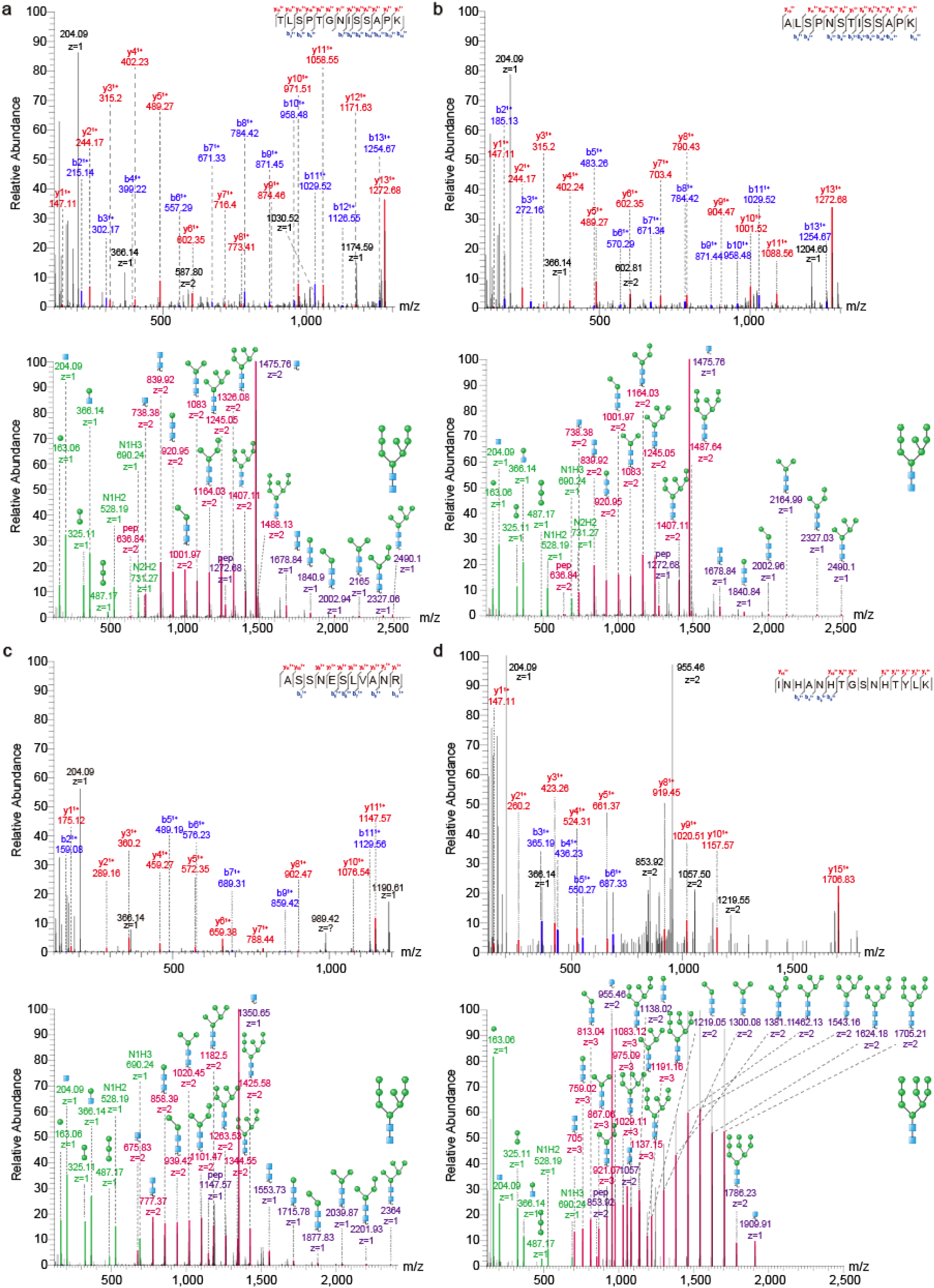
Intact glycopeptides of INM proteins. **a**-**d**, Intact glycopeptides of INM proteins SUN1 (**a**), SUN2 (**b**), TMPO (**c**) and LEMD3 (**d**) from mESCs.

**Extended Data Fig. 2.**
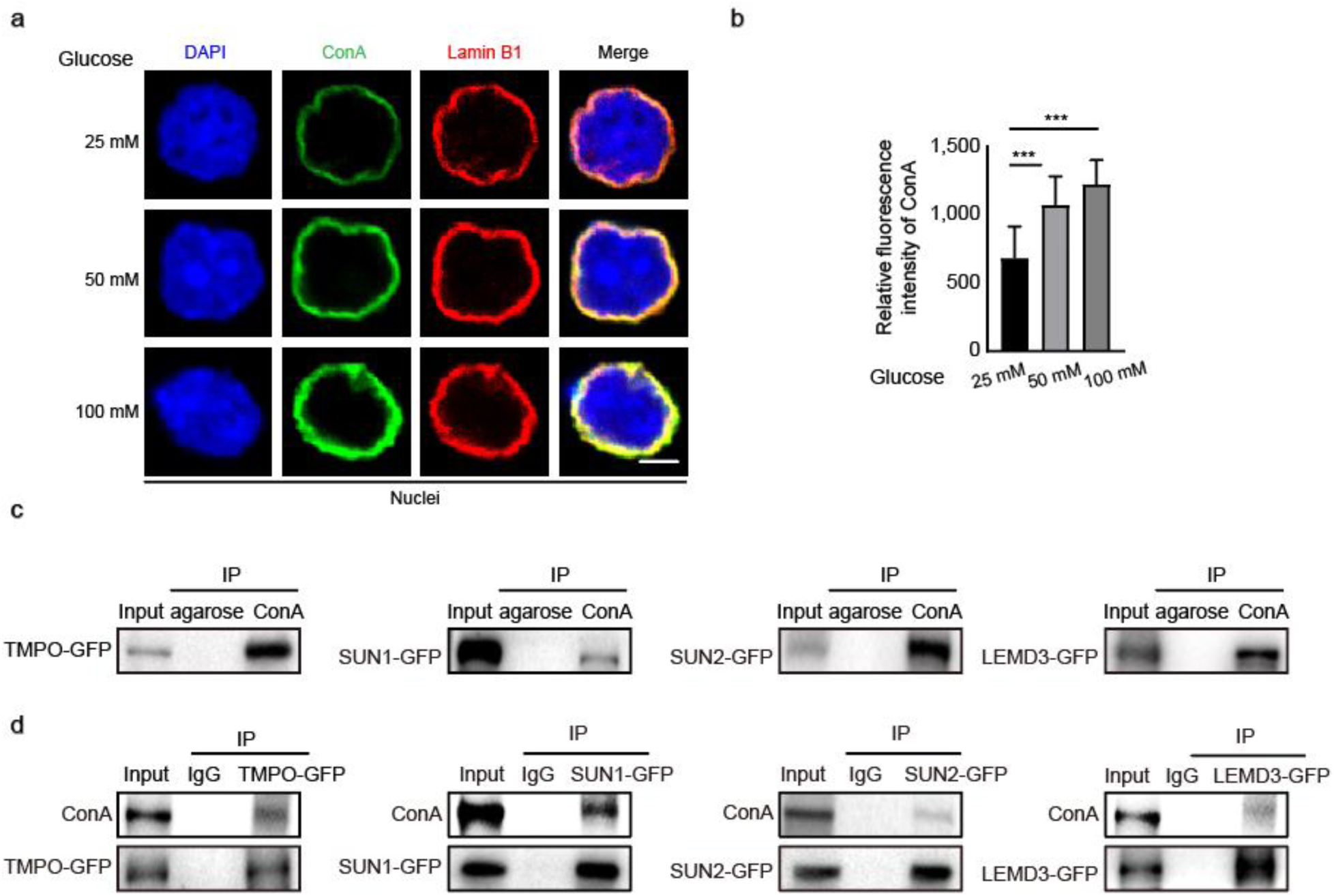
Validation of *N*-glycosylation of INM proteins. **a**, Immunofluorescence analysis of cell nuclei treated with different concentrations of glucose, scale bar is 5 μm. **b**, The quantitative analysis of immunofluorescence in **a**, data were presented as mean ± SD, *P*-values by two-tailed t-test, significance levels were indicated by asterisks: \**P* < 0.05; \*\**P* < 0.01; \*\*\**P* < 0.001. **c**, Blotting showing INM proteins in ConA-enriched *N*-glycosylated nuclear proteome, these experiments were performed three times, with similar results. **d**, Blotting of purified INM proteins to detect the *N*-glycosylated signals, these experiments were performed three times, with similar results.

**Extended Data Fig. 3.**
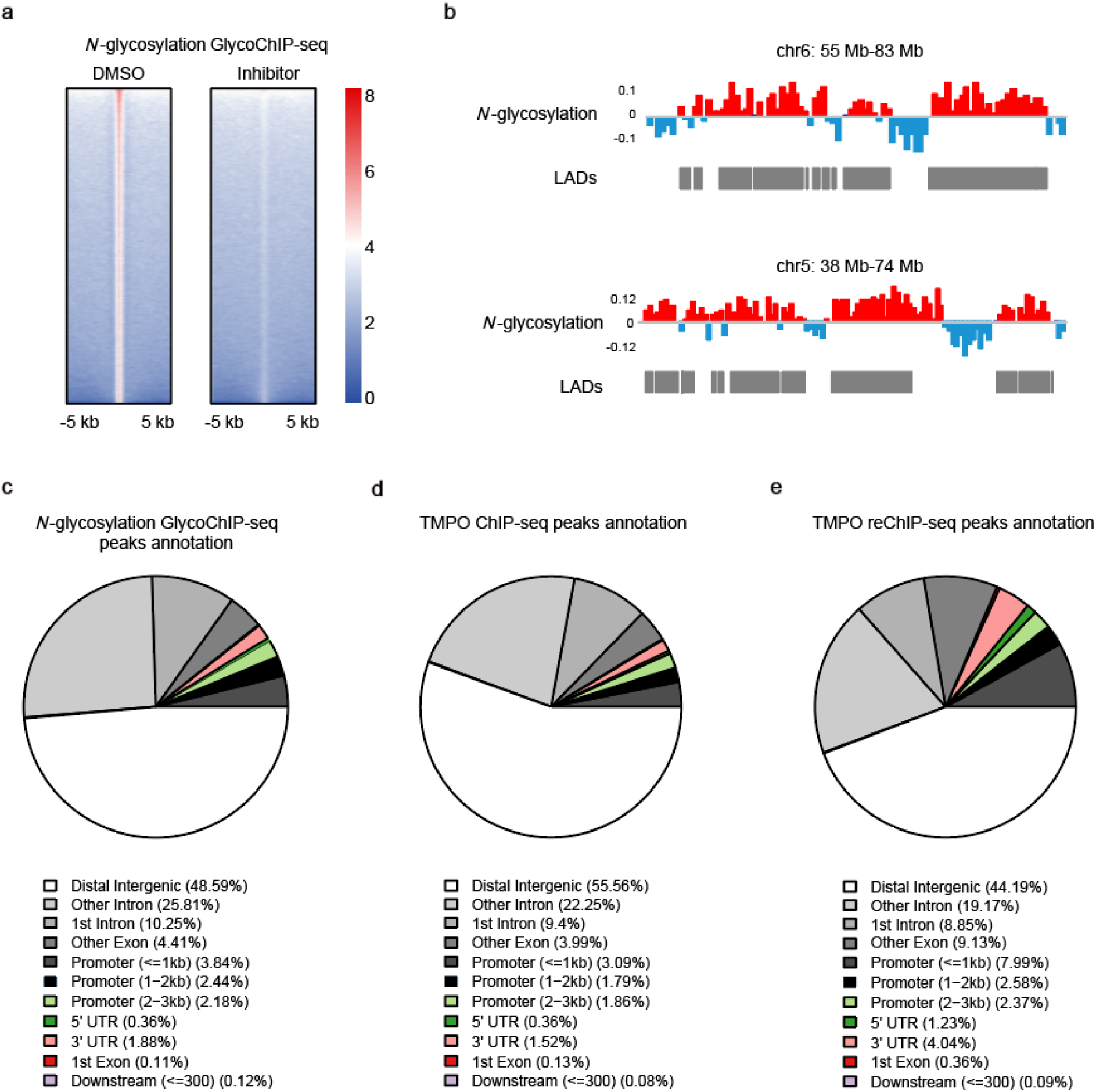
Bioinformatic analysis of ChIP-seq data. **a**, Heat maps illustrating the density of *N*-glycosylation GlycoChIP-seq reads before and after *N*-glycosylation inhibition in mESCs. **b**, Representative genome tracks of buffer-normalized *N*-glycosylation GlycoChIP-seq from mESCs. Gray bars indicate regions defined as LADs. **c**-**e**, The distribution of *N*-glycosylation GlycoChIP-seq peaks (**c**), TMPO ChIP-seq peaks (**d**), and *N*-glycosylated TMPO reChIP-seq peaks (**e**) with indicated genomic feature.

**Extended Data Fig. 4.**
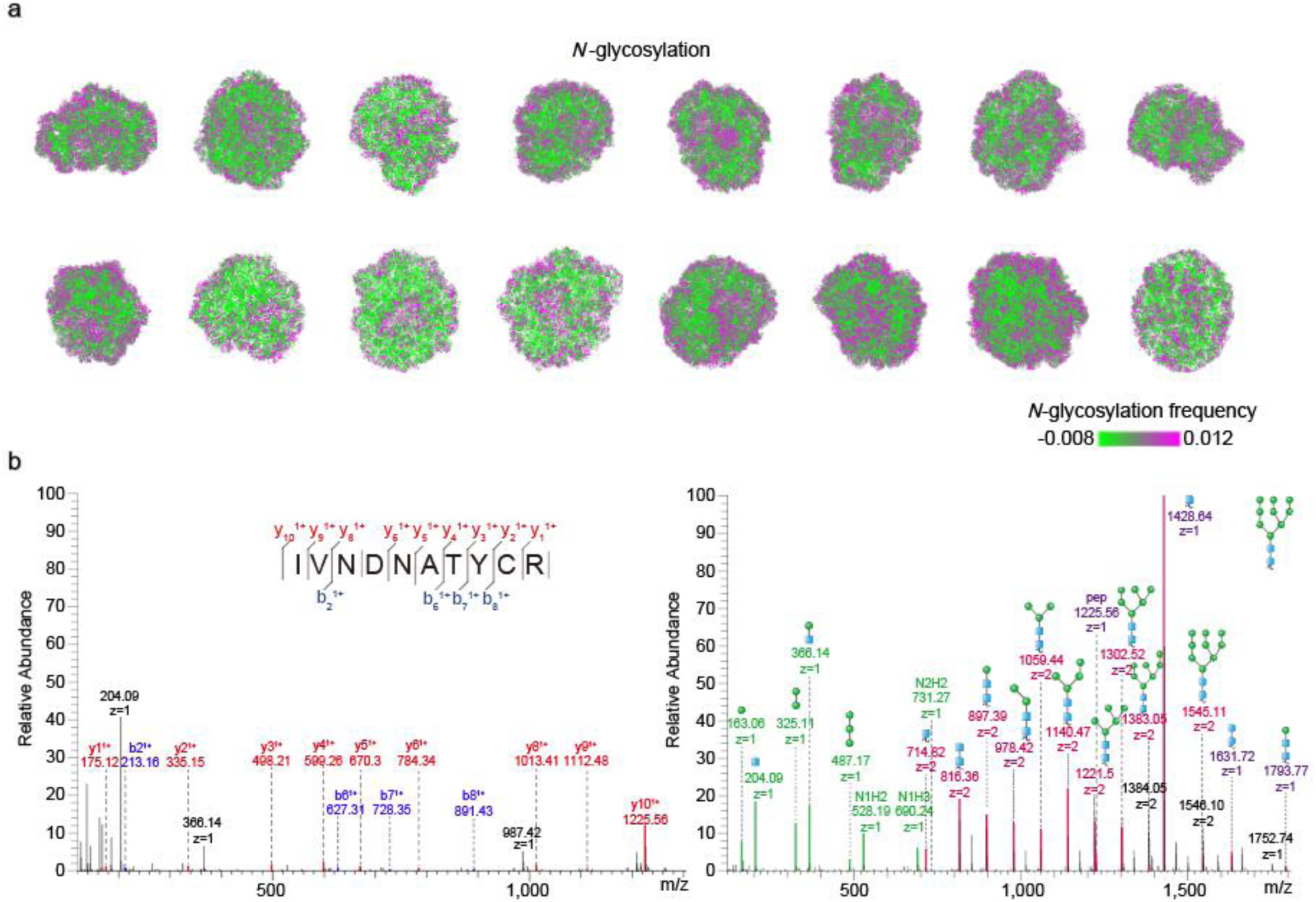
Representative 3D genome structure of mESCs and representative mass spectra of an intact glycopeptide identified from NOP56. **a**, Representative 3D genome structure of mESCs, colored by *N*-glycosylation frequency, cells were chosen randomly. **b**, Representative mass spectra of an intact glycopeptide identified from NOP56.

**Extended Data Fig. 5.**
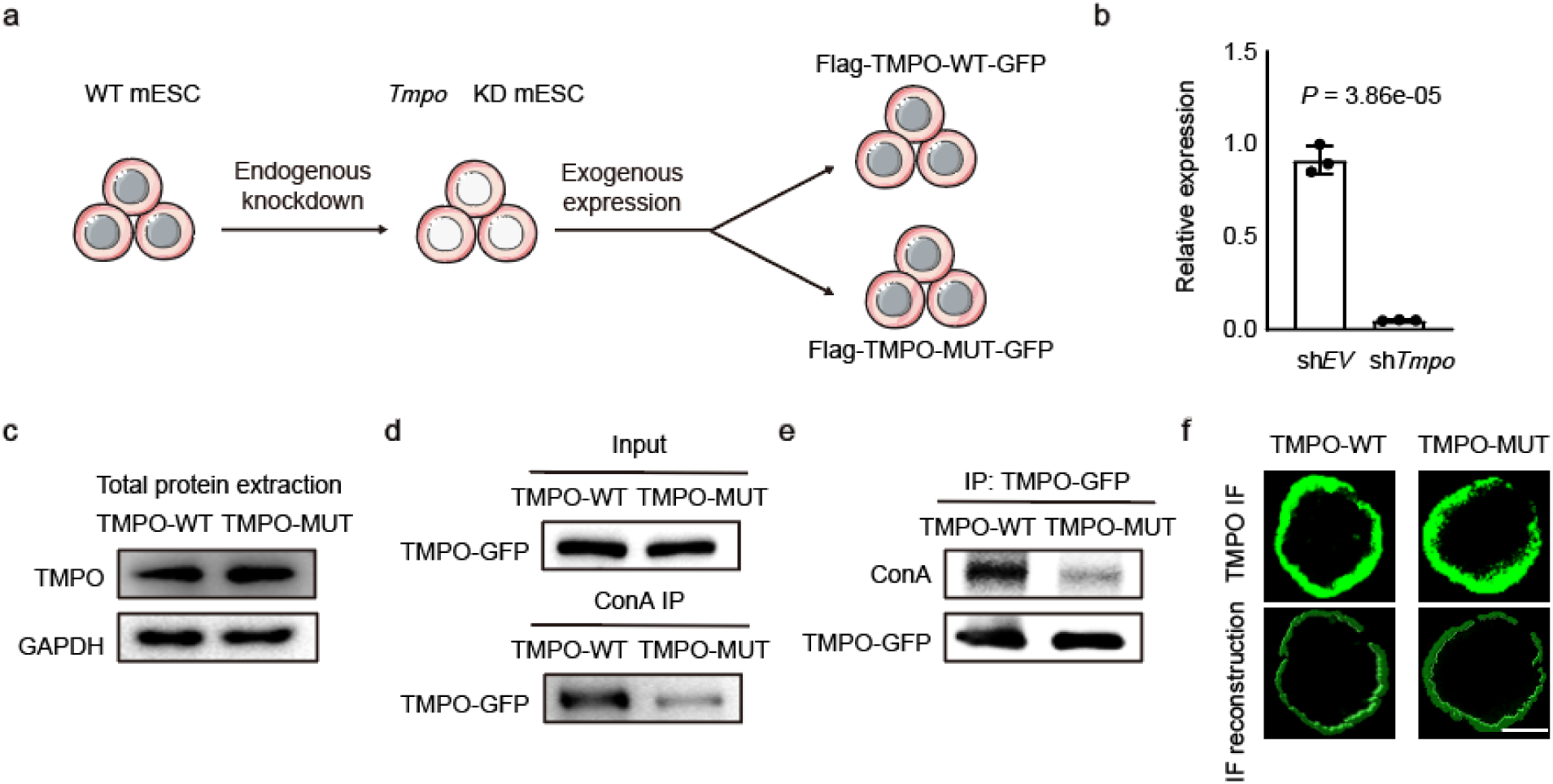
Construction and validation of the cell line with the TMPO *N*-glycosite mutation. **a**, Schematic diagram of constructing TMPO-WT and TMPO *N*-glycosite mutation (TMPO-MUT) cell lines. **b**, Quantitative PCR analysis of *Tmpo* expression after the knockdown of *Tmpo*, data were presented as mean ± SD (n = 3 biologically independent samples), *P*-value by two-tailed t-test. **c**, Blotting of total cell lysates from TMPO-WT cell line and TMPO-MUT cell line to detect the expression of TMPO, GAPDH was shown as loading control, these experiments were performed three times, with similar results. **d**, Blotting analysis to detect changes in the levels of TMPO in ConA-enriched *N*-glycosylated nuclear proteome, these experiments were performed three times, with similar results. **e**, Blotting of purified INM protein TMPO from TMPO-WT cell line and TMPO-MUT cell line to detect the *N*-glycosylated signals, these experiments were performed three times, with similar results. **f**, Immunofluorescence and reconstruction of TMPO in TMPO-WT cell line and TMPO-MUT cell line. Scale bar is 4 μm, this experiment was performed three times, with similar results.

**Extended Data Fig. 6.**
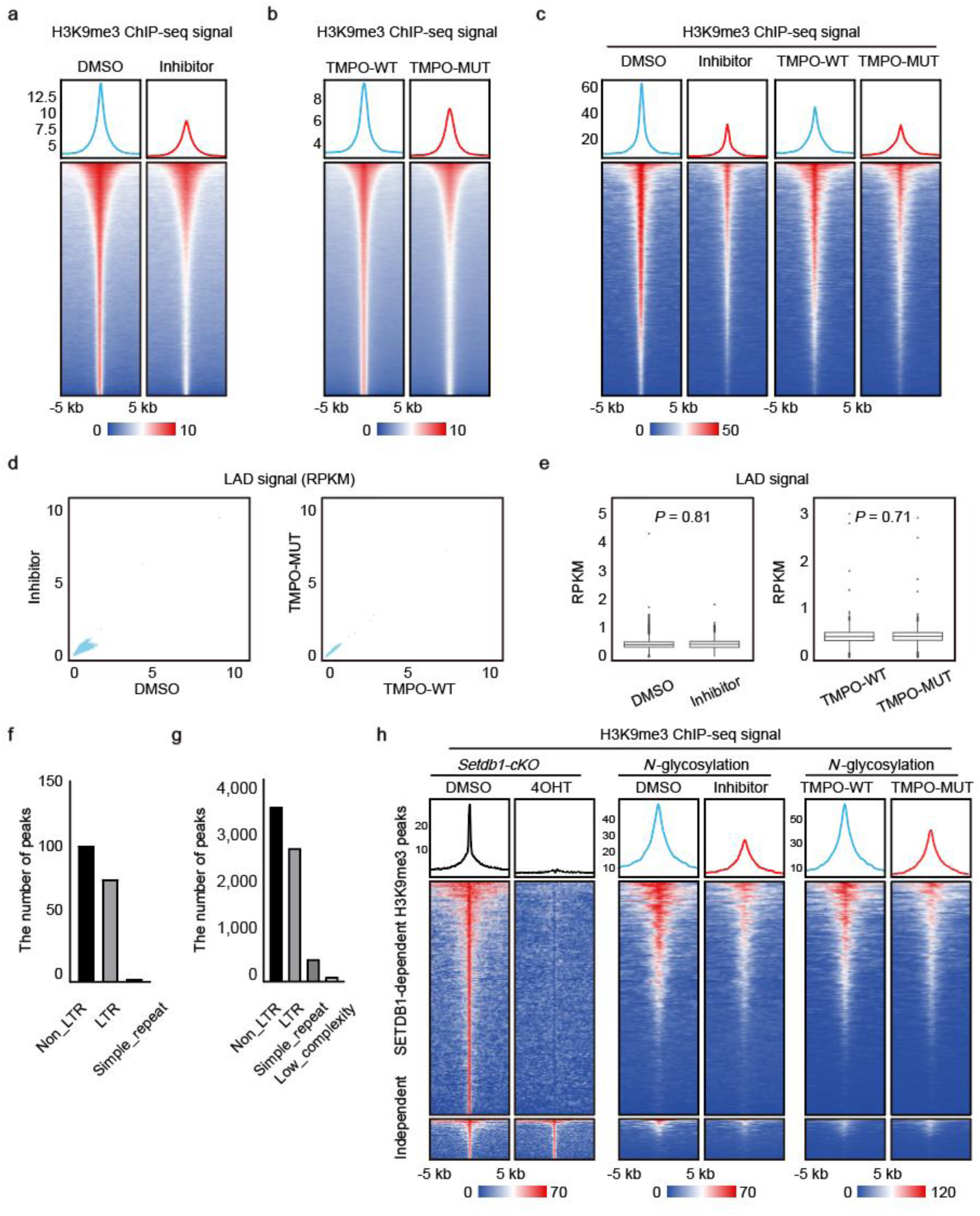
*N*-glycosylation inhibition and TMPO *N*-glycosite mutation both downregulate H3K9me3. **a**, Heat maps illustrating the density of H3K9me3 ChIP-seq reads before and after *N*-glycosylation inhibition. **b**, Heat maps illustrating the density of H3K9me3 ChIP-seq reads before and after TMPO *N*-glycosite mutation. **c**, Heat maps illustrating the density of H3K9me3 ChIP-seq reads on downregulated H3K9me3-binding peaks after *N*-glycosylation inhibition. **d**, Scatter plots showing overall LAD signals before and after *N*-glycosylation inhibition, and before and after TMPO *N*-glycosite mutation. **e**, Box plots showing signals of LAD before and after *N*-glycosylation inhibition, and before and after TMPO *N*-glycosite mutation, *P*-values by t-test. **f**, Bar graph showing the enriched regions of overlapping differential H3K9me3 peaks following *N*-glycosylation inhibition and TMPO *N*-glycosite mutation. **g**, Bar graph showing the enriched regions of overlapping peaks between *N*-glycosylation and H3K9me3. **h**, Heat maps illustrating the density of H3K9me3 ChIP-seq reads on SETDB1-dependent and SETDB1-independent H3K9me3-binding peaks, and read-count tag density pileups of H3K9me3 profiles on SETDB1-dependent H3K9me3 peaks.

**Extended Data Fig. 7.**
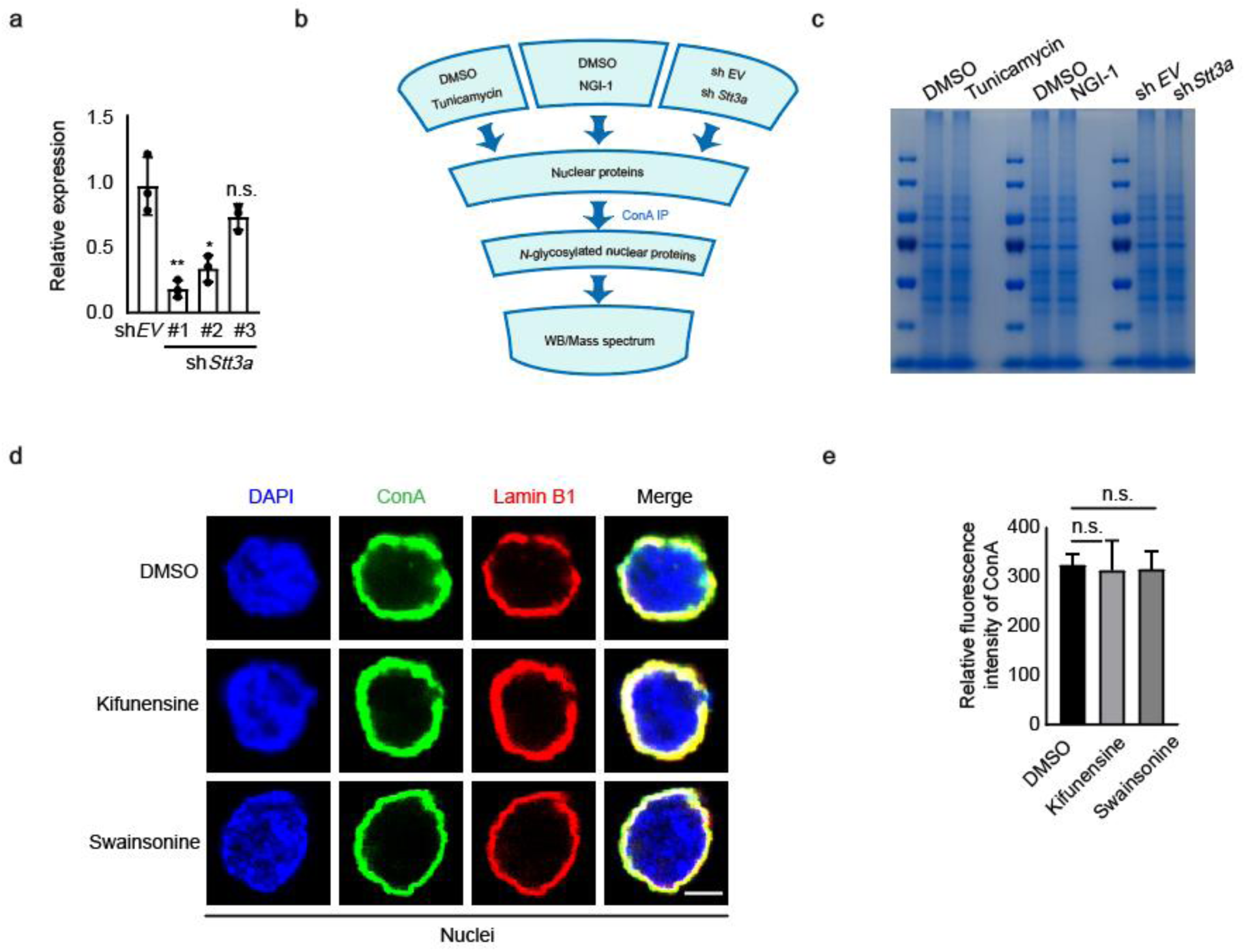
Canonical *N*-glycan biosynthetic machinery in ER contributes to the *N*-glycosylation of INM proteins. **a**, Quantitative PCR analysis of *Stt3a* expression after the knockdown of *Stt3a*, data were presented as mean ± SD (n = 3 biologically independent samples), *P*-values by two-tailed t-test, significance levels were indicated by asterisks: \**P* < 0.05; \*\**P* < 0.01; \*\*\**P* < 0.001 (not significant, denoted as n.s.). **b**, Schematic diagram of ConA-enriched *N*-glycosylated nuclear proteome from the cells subjected to the three different treatments. **c**, Coomassie blue staining of nuclear proteins used for ConA-enriched *N*-glycosylated nuclear proteome analysis, these experiments were performed three times, with similar results. **d**, Immunofluorescence of cell nuclei treated with kifunensine and swainsonine, scale bar is 5 μm. **e**, The quantitative analysis of immunofluorescence in **d**, data were presented as mean ± SD, *P*-values by two-tailed t-test, not significant, denoted as n.s..

**Extended Data Fig. 8.**
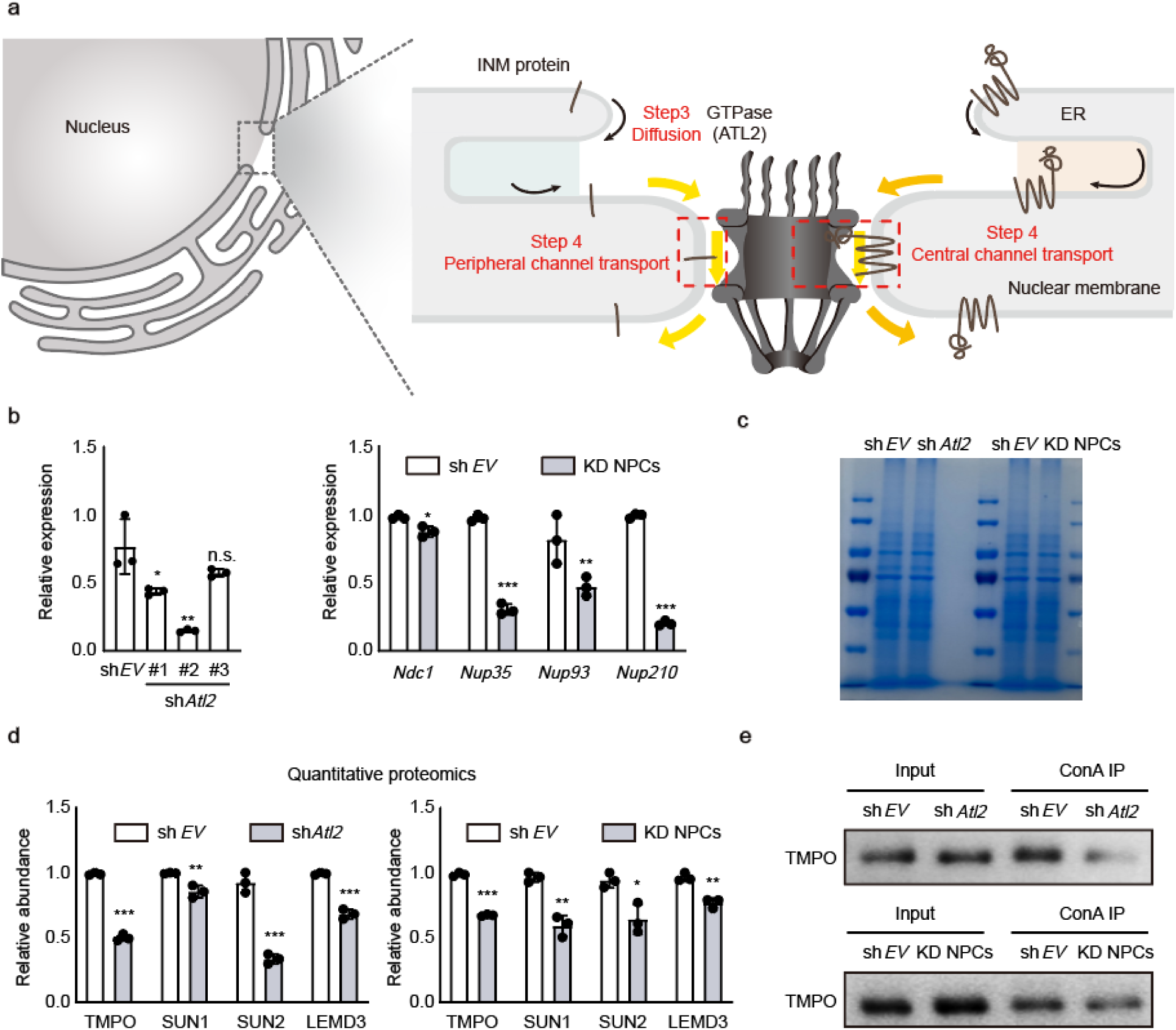
Knockdown of *Atl2* and nuclear pore complexes (NPCs) affect the *N*-glycosylation of INM proteins. **a**, An overview of the diffusion of INM proteins in the ER and their transport into nucleus. **b**, Quantitative PCR analysis of *Atl2* and NPCs expression, data were presented as mean ± SD (n = 3 biologically independent samples), *P*-values by two-tailed t-test, significance levels were indicated by asterisks: \**P* < 0.05; \*\**P* < 0.01; \*\*\**P* < 0.001 (not significant, denoted as n.s.). **c**, Coomassie blue staining of nuclear proteins used for ConA-enriched *N*-glycosylated nuclear proteome analysis, these experiments were performed three times, with similar results. **d**, Relative abundance of INM proteins in ConA-enriched *N*-glycosylated nuclear proteome detected by quantitative proteomics, data were presented as mean ± SD (n = 3 biologically independent samples), *P*-values by two-tailed t-test, significance levels were indicated by asterisks: \**P* < 0.05; \*\**P* < 0.01; \*\*\**P* < 0.001 (not significant, denoted as n.s.). **e**, Blotting analysis to detect changes in the levels of the INM protein TMPO in ConA-enriched *N*-glycosylated nuclear proteome, these experiments were performed three times, with similar results.

